# BioGCD: A Biology-Guided Multimodal Framework for Compound and Combination Discovery

**DOI:** 10.64898/2026.11.20.756733

**Authors:** Jian Wen, Zhihui Xu, Siqing Yue, Changzhen Weng, Ang Lin, Shentian Zhuang, Jinxian Wang, Xiaodan Fan, Xihan Zhang, Hanwei Wang, Kun Guo, Peidong Wu, Yubo Dong, Heqi Liu, Zutan Li, Weiwei Hu, Yong Yang, Cong Pian

## Abstract

Phenotype-based transcriptomic screening provides a powerful approach for therapeutic discovery by capturing how compounds reshape cellular states. However, existing perturbation models lack biological constraints, limiting generalizability, mechanistic interpretability, and the ability to prioritize experimentally validated candidates. Here, we developed BioGCD, a biological knowledge-guided multimodal framework that integrates four gene-level prior graphs and a compound-aware pathway-level prior with cellular states, molecular structures and dosage information to learn biologically informed perturbation patterns. BioGCD outperformed existing models in predicting compound induced transcriptomic responses across diverse cellular contexts, achieving a Pearson correlation coefficient of 0.912 between predicted and observed responses. Beyond prediction, the model enabled mechanistic interpretation of compounds effects by attributing predicted activities to specific molecular substructures and inferring potential pathways associated with compound responses. The proportion of the top 30 candidates supported by literature evidence or experimental validation varied across cell lines, reaching 70% in a melanoma cell line and a liver cancer cell line, and 30% in a breast cancer cell line. For previously uncharacterized agents, experimental screening of the top 50 candidates identified Z6412951658 as an anti-melanoma hit. Moreover, the BioGCD prediction-based model correctly classified 8 of 10 randomly selected compound combinations as synergistic or non-synergistic in melanoma cells. Overall, BioGCD establishes a biologically grounded AI framework for learning generalizable perturbation representations, enabling systematic therapeutic discovery through accurate prediction and biological insight.

## Introduction

Phenotype-based screening aims to identify compounds that modulate disease-relevant cellular states, providing a powerful approach for drug discovery and repositioning. With the development of large-scale transcriptomic technologies, gene expression profiles have become a key molecular representation of cellular states and disease phenotypes. By projecting disease- and compound-induced signatures into a shared transcriptional space, transcriptional profiling enables systematic comparison across diseases, compounds and biological pathways. In this framework, similar signatures may indicate shared mechanisms of action, whereas compounds that reverse disease-associated transcriptional programs may have therapeutic potential.

The Connectivity Map established this transcriptome-based paradigm by linking small-molecule perturbations with genome-wide expression profiles in human cell lines^1^. Subsequent studies demonstrated that transcriptional response similarity and network perturbation patterns can support mechanism-of-action prediction and drug repositioning^2,3^, while disease-drug signature reversal has been used to identify therapeutic candidates across diverse disease contexts^4–7^. The L1000-based next-generation Connectivity Map substantially expanded perturbational profiling to more than one million chemical and genetic perturbation profiles^8,9^. However, even resources of this scale cover only a small portion of the combinatorial space defined by compounds, cell types, disease states, doses and treatment conditions. The challenge has motivated the development of computational methods that leverage deep learning and large-scale perturbational resources to predict transcriptional responses.

Early counterfactual models, such as scGen, showed that perturbation-induced state transitions can be learned as latent space shifts across cell types and conditions^10^, but did not explicitly model compound identity. Subsequent compound-focused methods incorporated chemical dosage and cellular context to predict compound-induced transcriptional changes, including DLEPS, DeepCE, MultiDCP and CIGER^11–14^. TranSiGen and PRnet further shifted from fixed perturbational signatures toward basal state conditioned response prediction, using baseline expression states and molecular representations to infer chemical-induced transcriptional changes in unobserved contexts^15,16^. Many of these models remain constrained by dependence on training cell lines or fixed dose and exposure-time settings, whereas others predict only a limited subset of genes, restricting their ability to comprehensively characterize compound-induced transcriptomic responses. In parallel, studies using single-cell perturbation datasets have addressed cellular heterogeneity in perturbation responses. A major line of work decomposes expression variation into basal state, perturbation, dose and covariate effects, enabling counterfactual prediction for unseen compounds, doses or combinations^17–20^. Another complementary line treats perturbation response as a distributional mapping problem, using optimal transport to infer transitions from untreated to perturbed cellular states, as in CellOT and CINEMA-OT^21,22^.

Despite recent progress, most existing models rely primarily on data-driven representations with limited incorporation of biological regulatory priors, which constrains their ability to capture mechanistically meaningful perturbation effects and generalize to unseen cellular or chemical contexts. This limitation is particularly critical for predicting compound induced transcriptional responses, because reliable response prediction forms the basis for comparing compound-induced signatures with disease-associated signatures and quantifying the extent to which a compound may reverse disease-related transcriptional dysregulation. To address these challenges, we developed BioGCD, a biological knowledge-guided multimodal framework that integrates cellular states, molecular structures, dosage information, four gene-level prior graphs, and a compound-aware pathway-level prior to learn context-specific perturbation patterns and rank compounds by reversal potential. BioGCD achieved superior unseen-durg response prediction against competing models, and enabled multilevel interpretation by identifying activity-related molecular substructures and cancer-relevant signatures. Extending these capabilities, the framework prioritized therapeutically relevant candidates across multiple cancer types, with many top-ranked candidates supported by prior evidence or experimental validation. Experimental evaluation further identified Z6412951658 as a potent anti-melanoma hit among previously uncharacterized melanoma candidates and confirmed the synergistic activity of predicted compound combinations in melanoma cells. Collectively, this study establishes a biologically grounded AI framework for systematic drug repurposing and discovery, as well as rational recommendation of synergistic compound combinations.

## Results

### Overview of the BioGCD framework for compound recommendation

The L1000 resource contains a large number of compound-induced gene-expression profiles, but these data are generated under diverse experimental conditions and include replicate measurements with variable signal quality. To obtain a standardized dataset suitable for model development, we retained L1000 profiles generated after 24h of compound exposure^9^ (Fig. 1a). Replicate profiles corresponding to the same perturbation condition were averaged before quality control. Low-variance and outlier profiles were subsequently removed. The curated dataset comprised more than 100,596 perturbation profiles and 164 control profiles across 164 cell lines.

**Figure 1.**
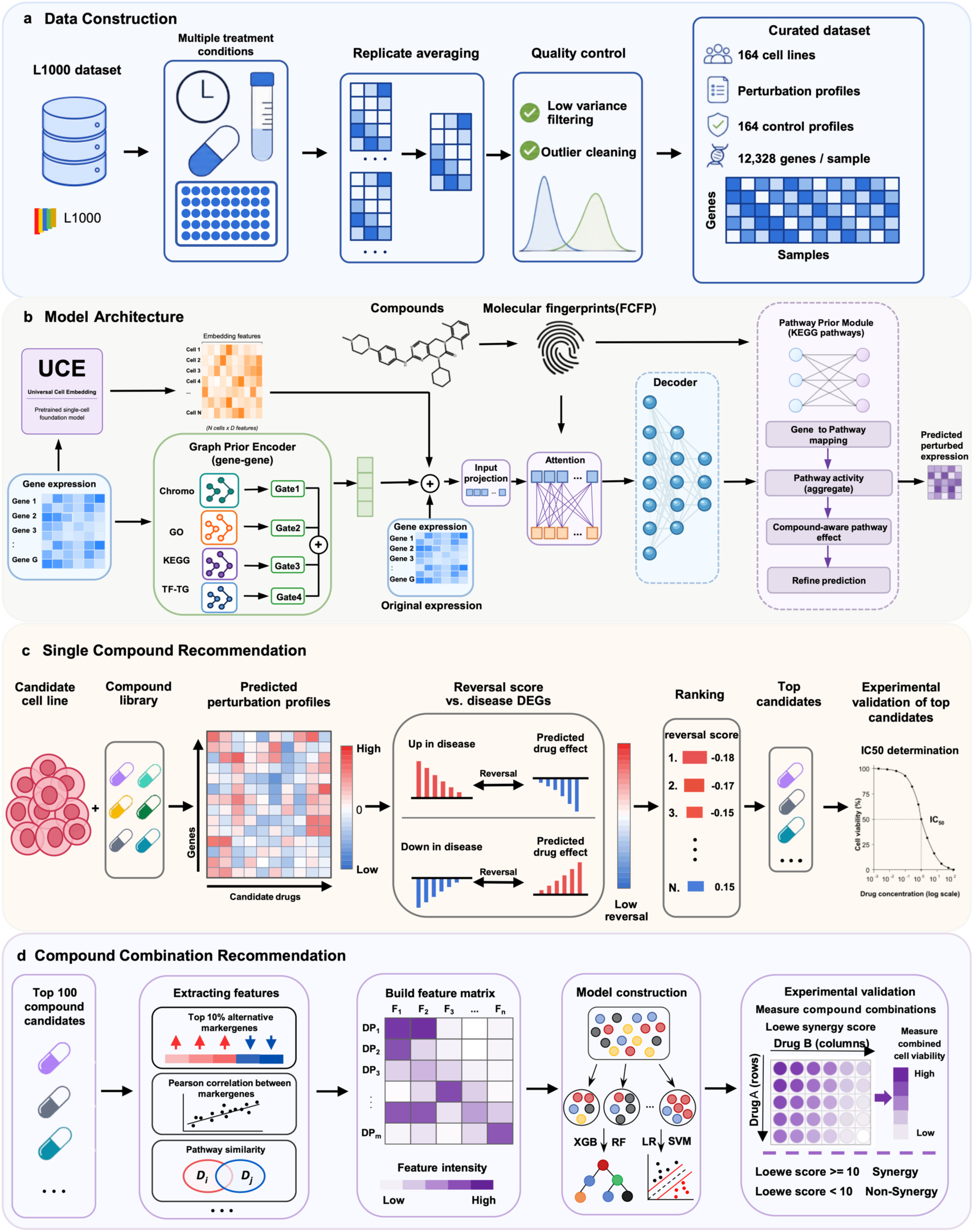
Overview of BioGCD framework for compound response prediction and prioritization. (a) Construction and preprocessing of the L1000 perturbation dataset. (b) BioGCD architecture for predicting compound induced transcriptional responses. (c) Single compound prioritization based on disease signature reversal and experimental validation. (d) Combination synergy prediction followed by experimental validation using Loewe synergy scoring.

BioGCD predicts gene expression responses to unseen compounds by jointly integrating cellular context, compound structure, dose, and biological prior knowledge (Fig. 1b). To capture cellular state, baseline gene expression profiles are passed through a pretrained foundation model, Universal Cell Embedding (UCE), which produces a 1,280-dimensional representation that summarizes transcriptomic context across diverse cell states^23^. Candidate compounds are encoded using FCFP4 molecular fingerprints^24^, and compound dose is projected into the shared latent space by a dose encoder. Biological priors, including chromosomal proximity, Gene Ontology, KEGG pathways, and transcription factor-target regulation, are represented as gene-level graphs and encoded to capture prior-informed gene dependencies^25,26^. These graph-derived features are fused with UCE-based cellular representations and baseline expression features, and the resulting cellular features are integrated with molecular features through cross-attention over latent tokens^27^. Finally, BioGCD decodes the integrated representation into compound-induced expression changes, which are added to the baseline profile through a residual connection, while a KEGG pathway-prior module further refines gene-level predictions by modeling compound-aware pathway activities.

To translate predicted transcriptional responses into therapeutic hypotheses, we designed a two-stage prioritization workflow for single-compound and combination-compound discovery. For single-compound prioritization, a target cell line and a candidate compound library were first specified (Fig. 1c). BioGCD then predicted the post-treatment expression profile of the selected cell line for each candidate compound, and log₂ fold-change values were calculated relative to the corresponding control profile. Each predicted response was compared with the upregulated and downregulated disease signatures using a rank-based reversal score adapted from the Connectivity Map framework^1^. Lower reversal scores indicate a stronger predicted ability to reverse the disease-associated up- and downregulated transcriptional signatures. Compounds were ranked in ascending order of reversal score, and the top 30 candidates were selected for experimental validation using cell viability and half-maximal inhibitory concentration assays.

For combination-compound recommendation, we developed a BioGCD-based workflow that uses predicted transcriptional and pathway-level responses to characterize candidate two-compound combinations (Fig. 1d). For each candidate compound pair, BioGCD first predicted the gene-expression response induced by each single agent in the target cell line. We then computed multiple pairwise features, including the overlap ratio of the most strongly perturbed genes, the Pearson correlation between predicted compound induced expression changes, and the Jensen-Shannon similarity between pathway-level response profiles generated by the KEGG pathway-prior module. Because these features showed cell-line dependent differences between synergistic and non-synergistic pairs, we further expanded them into a broader feature set and trained machine-learning classifiers to evaluate their ability to predict compound-pair synergy. Candidate combinations were randomly sampled across the resulting feature space to ensure coverage of compound pairs with diverse transcriptional similarity and response-overlap patterns. Selected combinations were subsequently tested using cell viability assays and Loewe synergy scoring.

### BioGCD outperforms existing models in unseen compounds response prediction

To evaluate the generalization capability of BioGCD to unseen compounds, we split the dataset into training, validation, and test sets at an 8:1:1 ratio (Fig. 2a). Model training, validation-based model selection, and test-set evaluation were repeated using three random seeds. Performance was assessed by comparing the predicted perturbation responses with the corresponding ground truth responses in the test set. chemCPA^17^, CPA^18^, PRnet^16^, and TranSiGen^15^ were evaluated on the same preprocessed dataset using identical compound-disjoint splits and evaluation procedures to ensure a fair comparison.

**Figure 2.**
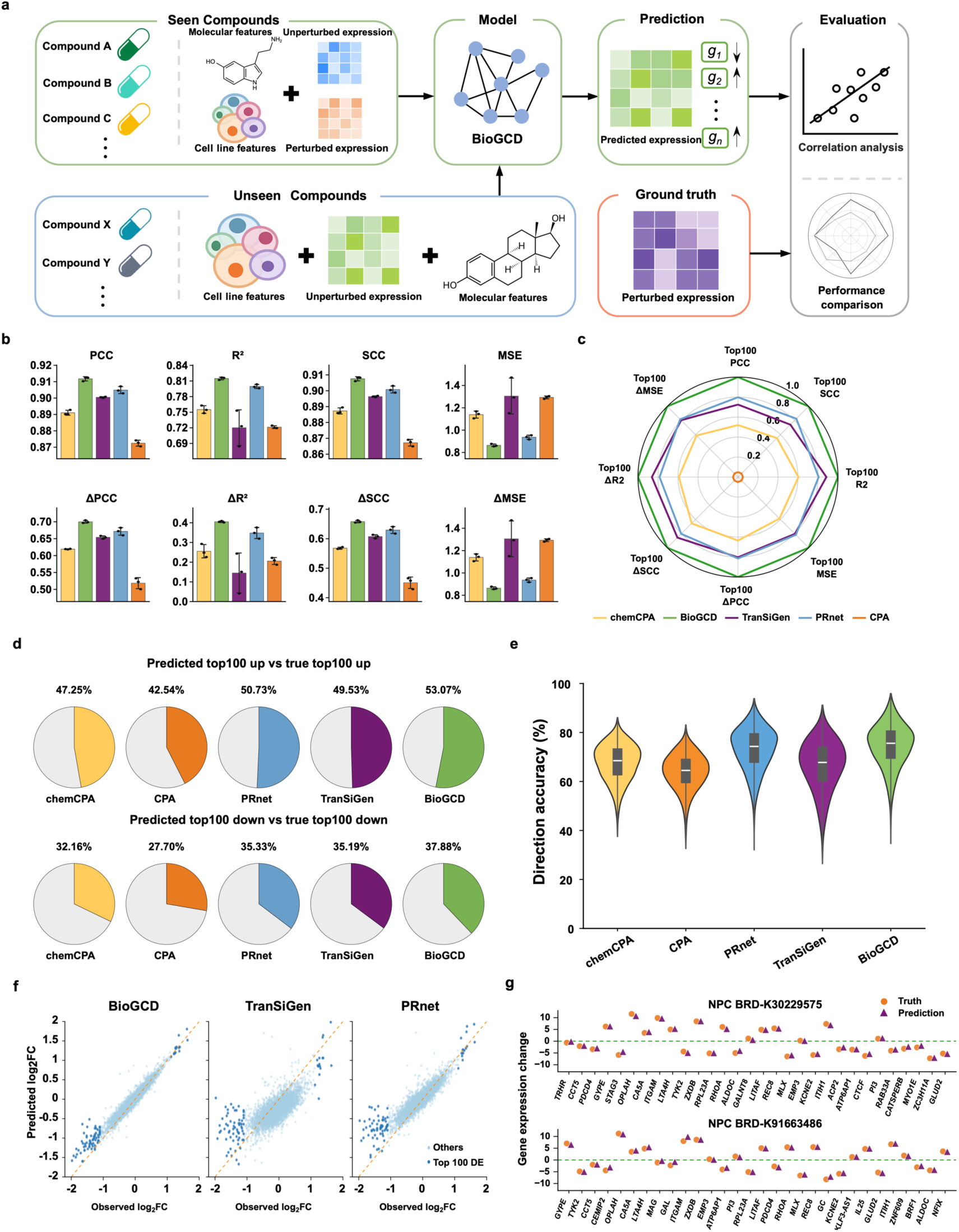
Evaluation of BioGCD’s generalization to unseen compounds across multiple criteria. (a) Compound-disjoint evaluation design for unseen-compound prediction. Compounds are divided into training, validation and test sets at an 8:1:1 ratio, keeping all profiles for each compound within one subset. Predictions for held-out compounds are compared with experimentally measured perturbed expression profiles. (b) Overall performance comparison across eight evaluation metrics. Δ represents expression changes relative to matched controls. Error bars represent the standard deviation across three independent splits. (c) Predictive performance for the top 100 differentially expressed genes. Mean metric values are min–max scaled across models. (d) Overlap between predicted and observed top 100 upregulated and downregulated genes. Colored sectors and percentages represent the proportion of the observed gene set recovered by each model. (e) Distributions of directional accuracy across all combinations of cell lines and compounds. Directional accuracy is the proportion of genes for which predicted and observed expression changes relative to matched controls have the same direction. (f) Comparison of predicted and observed log₂ fold changes under a representative perturbation condition. Dark blue points indicate the 100 most strongly perturbed genes, whereas light blue points represent all remaining genes. (g) Representative comparisons between predicted and observed gene-expression. Orange circles denote observed values, purple triangles denote predictions, and dashed horizontal lines indicate zero change.

Under this evaluation framework, BioGCD achieved the best mean performance among all five models across both global expression and perturbation response metrics (Fig. 2b). For global expression prediction, it obtained a PCC of 0.912 ± 0.001, SCC of 0.908 ± 0.001, R² of 0.815 ± 0.003, and MSE of 0.864 ± 0.012, while also ranking highest in ΔPCC, ΔSCC, and ΔR² for perturbation-induced expression changes. Consistently, the advantage of BioGCD was retained on the top 100 differentially expressed genes (Fig. 2c), with the highest overlap observed for the true top-100 upregulated and downregulated genes at 53.07% and 37.88%, respectively (Fig. 2d). Moreover, the distribution of direction accuracy further confirmed the superior performance of our model in predicting the direction of gene expression changes (Fig. 2e).

We also assessed whether BioGCD could generalize to perturbations induced by different compound concentrations by comparing its performance with CPA, chemCPA and PRnet across all genes and the top 100 most perturbed genes. BioGCD achieved the best aggregate performance in both evaluations, yielding the highest PCC, SCC and R^2^ values and the lowest MSE among all methods (Table S1). Dose-stratified analyses further confirmed that BioGCD maintained consistently high accuracy across all concentration intervals, including for strongly responsive genes represented by the top 100 most perturbed genes. t-SNE visualization of all perturbation profiles revealed a structured transcriptional landscape, in which profiles formed distinct cell-line-specific manifolds with clear dose-dependent gradients (Fig. S1a). Independent t-SNE analyses of representative cell lines further showed dose-dependent separation of transcriptional profiles induced by 516 compounds across five dose levels within each cellular context (Fig. S1b).

At the transcriptome-wide level, BioGCD also better captured global perturbation properties. Across cell lines, it showed the most consistent agreement with true perturbation responses (Fig. S2a). For all evaluated models, discrepancies from the true distributions of transcriptome-wide impact and effective DEG numbers were observed (Fig. S2b). Nevertheless, PRnet, TranSiGen, and BioGCD produced distributions relatively closer to the ground truth. Among them, BioGCD captured stronger transcriptome-wide perturbation effects, thereby better recovering large-magnitude gene expression changes.

To assess prediction fidelity at the gene level, the agreement between predicted and observed gene expression changes after perturbation was analyzed. A closer match to the measured responses was observed for BioGCD, with predicted changes more tightly distributed along the diagonal than those of competing methods (Fig. 2f). Its ability to capture condition-specific gene-level responses to perturbation was further confirmed by representative unseen-compound examples (Fig. 2g). Gene quantile analysis showed that prediction errors increased for strongly perturbed genes, but stable and competitive performance was maintained by BioGCD across response-strength groups, with high correlations and direction accuracy (Fig. S3).

### BioGCD captures interpretable representations for compound response prediction

The ability of BioGCD to learn biologically meaningful representations from multimodal input data was evaluated by analyzing the resulting latent space across major cellular contexts. For the ten cell types with the largest sample sizes, representations based on the original UCE and expression features showed partial organization by cell type, whereas the projected representations exhibited more distinct cell type dependent clustering (Fig. 3a).

**Figure 3.**
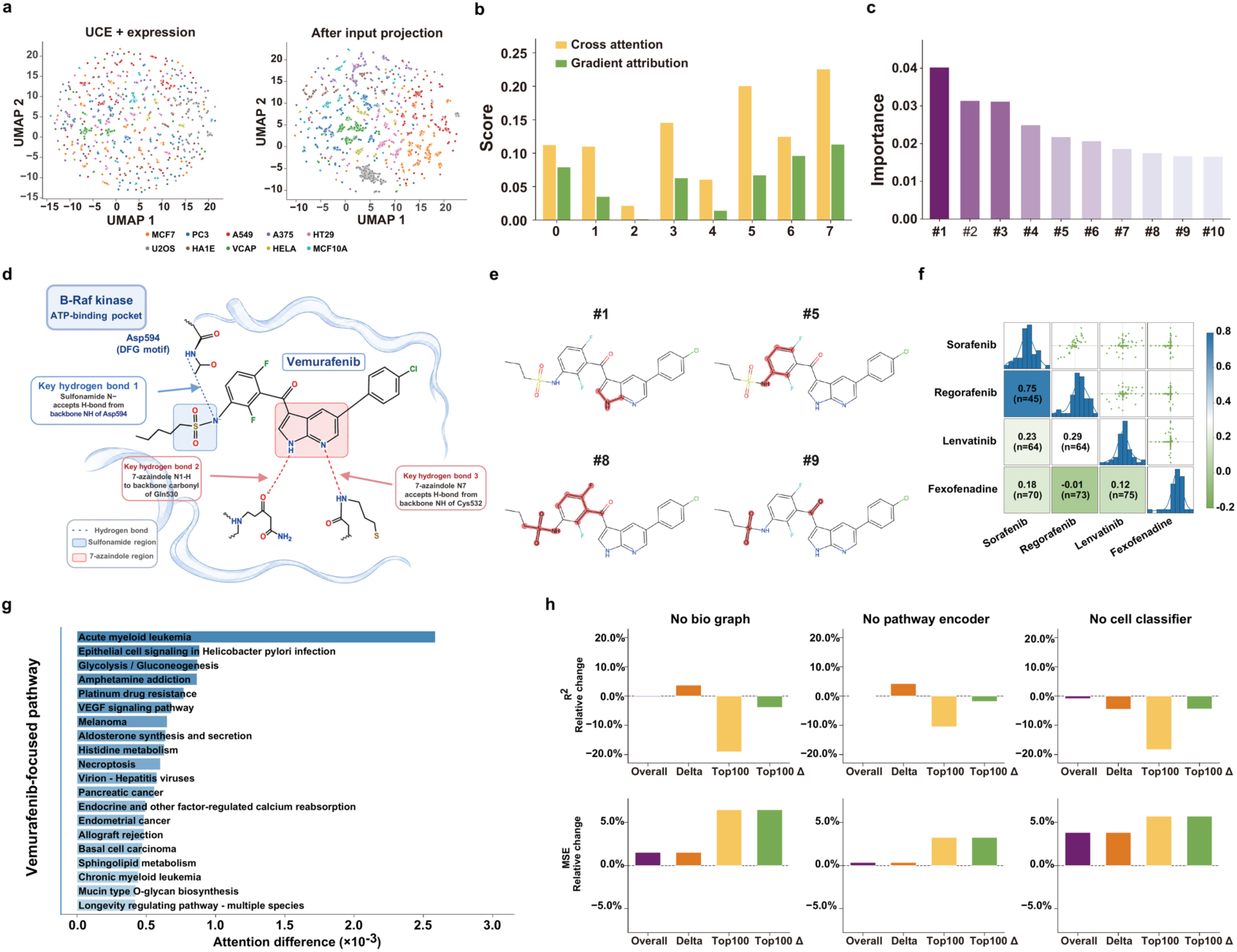
Multi-level interpretation of BioGCD predictions and assessment of individual model components. (a) UMAP visualization of cell-type organization before and after input projection. Colors distinguish the ten cell lines with the largest sample sizes. (b) Cross-attention and gradient attribution scores for vemurafenib tokens. The 1,024-bit FCFP4 is divided into eight consecutive 128-bit tokens, indexed from 0 to 7. (c) Importance scores for the ten highest ranked individual fingerprint bits of vemurafenib. Labels #1-#10 denote their positions in the ranked list. (d) Knowledge-guided schematic of vemurafenib-BRAF interactions within the ATP-binding pocket. The sulfonamide and 7-azaindole regions and their annotated hydrogen bonds with Asp594, Gln530, and Cys532 are highlighted. (e) Molecular substructures associated with representative fingerprint bits of vemurafenib with high attribution scores. Highlighted regions show where these fragments map onto the molecular structure. (f) Pairwise comparison of molecular attribution scores for sorafenib, regorafenib, lenvatinib, and fexofenadine. Each comparison includes the union of active FCFP4 bits in the two compounds. n indicates the number of included bits. (g) The top 20 KEGG pathways ranked by model derived attention differences. (h) Ablation study of the biological graph-prior encoder, pathway encoder, and cell classification module. Bars show percentage changes relative to the complete model.

We selected vemurafenib as a representative compound with well-characterized structure activity relationships to evaluate whether BioGCD attribution could recover activity-relevant molecular features. We performed attribution analysis for vemurafenib at both the token and individual-bit levels. Its 1,024-bit FCFP4 fingerprint was divided into eight consecutive 128-bit tokens for token-level interpretability analysis. Token 7 showed the strongest contribution based on both cross-attention and input-gradient product attribution, followed by tokens 5 and 6, which were also consistently highlighted by both methods (Fig. 3b). At the individual-bit level, bits 1010, 789, 498, and 174 were identified as the most influential fingerprint features for vemurafenib (Fig. 3c, Table S2).

Previous studies have shown that vemurafenib inhibits BRAF^V600E^ through critical hydrogen-bond interactions involving the sulfonamide and 7-azaindole regions. These include an interaction between the sulfonamide nitrogen and the backbone NH of Asp594 within the DFG motif, a hydrogen bond between the 7-azaindole N1-H and the backbone carbonyl of Gln530, and hydrogen-bond acceptance by the 7-azaindole N7 from the backbone NH of Cys532 (Fig. 3d)^28,29^. Notably, most of the top-ranked fingerprint bits that mapped to activity-related structural regions were located within the highly attributed tokens (Table S2). Structural interpretation of these fingerprint bits revealed that several corresponded to substructures involved in, or adjacent to, the reported hydrogen-bonding pharmacophore. Bit #1 primarily mapped to the nitrogen-containing heteroaromatic core, including the 7-azaindole-related fragment. Bits #5 and #8 highlighted the halogenated aromatic region and the adjacent sulfonamide/carbonyl environment, whereas bit #9 mapped to the sulfonamide- and carbonyl-associated region (Fig. 3e). Additionally, we assessed the consistency of molecular attribution patterns across the other four compounds. Sorafenib and regorafenib showed the highest FCFP4 fingerprint similarity and shared closely related chemical scaffolds, whereas lenvatinib and fexofenadine were structurally more distinct (Fig. S4a). We therefore calculated the Jaccard index of active FCFP4 bits and pairwise Spearman correlations between signed input-gradient product attribution scores over the union of active bits for each compound pair (Fig. 3f and Fig. S4b). Sorafenib and regorafenib exhibited the strongest attribution agreement, whereas correlations involving lenvatinib or the control compound fexofenadine were substantially lower. These analyses supported that BioGCD learned structure-dependent local attribution patterns.

A key innovation of our model is the KEGG pathway-prior module, which aggregates baseline gene expression into pathway-level activities and enables compound-specific attention over KEGG pathways. We next examined the pathway-attention profile of vemurafenib to assess whether the model highlighted biological processes relevant to its pharmacological activity. The top-ranked pathways included melanoma and glycolysis/gluconeogenesis (Fig. 3g). This pattern was consistent with vemurafenib’s established activity in BRAF-mutant melanoma^28^ and the role of glycolytic regulation in responses to BRAF inhibition^30^.

After confirming that BioGCD learned biologically meaningful global representations, structure dependent molecular attribution patterns, and mechanism associated pathway-attention profiles, we sought to determine which modules were essential for model performance. Each component, including the biological prior graph, pathway encoder, and cell-classification task module was removed individually and evaluated by ablation analysis (Fig. 3h). Although the relative global performance changed only modestly, top100 gene prediction was more strongly affected, with R^2^ decreasing by 19.07%, 10.51%, and 18.29%, respectively, and similar reductions were also observed for top100 delta R^2^. Slight increases in delta R^2^ in some ablations were accompanied by lower correlations and higher MSE, suggesting metric trade-offs rather than true performance improvements.

### BioGCD identifies diverse anti-cancer candidates

Using the melanoma cell line, we systematically prioritized single-agent candidates by applying BioGCD to valid compounds from the TargetMol T001 screening-compound database and predicting their transcriptional responses at 10 μM. The predicted responses were then compared with cancer associated gene expression signatures using a rank-based reversal score, with lower scores indicating stronger predicted reversal of the melanoma associated transcriptomic signature. Candidate compounds were visualized in both the predicted log₂ fold-change response space and the FCFP4 chemical-structure space (Fig. 4a, Table S3). The top 30 prioritized compounds clustered within a restricted region of the predicted response space, whereas they were more broadly distributed in the chemical-structure space. This pattern indicates that these compounds elicited similar predicted transcriptomic reversal effects despite exhibiting diverse chemical structures. Structure-based drug family assignment showed that they represented multiple chemically distinct families, including camptothecin analogues, CDK-like kinase inhibitors, anthracycline analogues, and FAK/PYK2 inhibitors (Fig. 4a). These findings indicate that BioGCD can identify structurally diverse compounds with convergent melanoma-reversing transcriptional effects.

**Figure 4.**
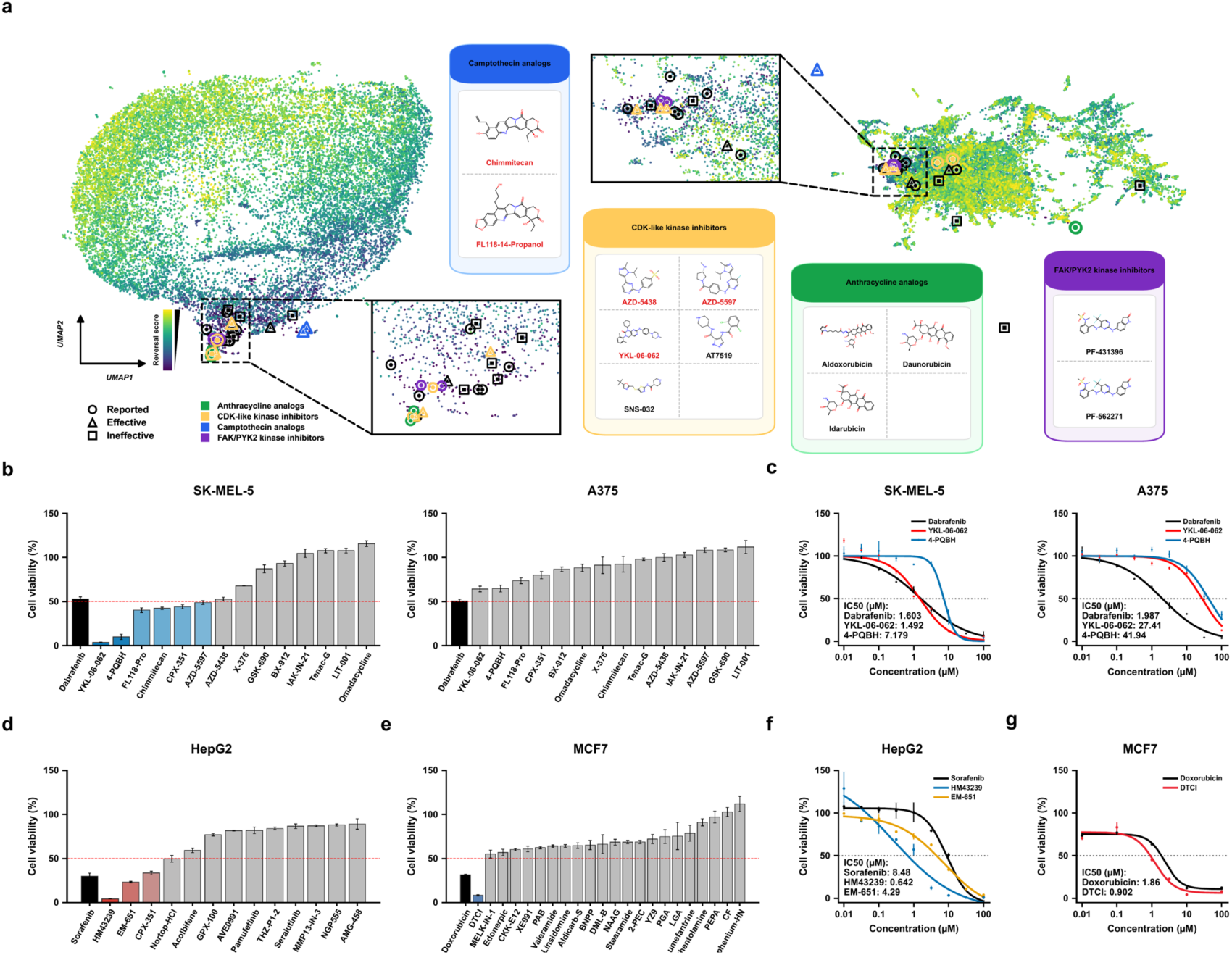
BioGCD prioritizes individual drug candidates with experimentally observed activity across cancer cell lines. (a) UMAP projections of candidate compounds in predicted transcriptional response space (left) and FCFP4 molecular fingerprint space (right). Point colors indicate reversal scores. Circles, triangles, and squares denote previously reported, experimentally effective, and experimentally ineffective compounds, respectively. Insets magnify regions containing candidate compounds and show representative molecular structures. (b) Cell viability screening of prioritized candidates in A375 and SK-MEL-5 cells, with dabrafenib as the reference drug. Colored and gray bars indicate mean viability below and at or above 50%, respectively. The dashed red line marks the 50% viability threshold. (c) Dose response curves for dabrafenib, YKL-06-062 and 4-PQBH in the two melanoma cell lines. (d) Cell viability screening of prioritized candidates in HepG2 hepatocellular carcinoma cells, with sorafenib as the reference compound. (e) Cell viability screening of prioritized candidates in MCF7 breast cancer cells, with doxorubicin as the reference drug. (f) Dose response curves for sorafenib, HM43239 and EM-651 in HepG2 cells. (g) Dose response curves for doxorubicin and 3,3-diethylthiacarbocyanine iodide in MCF7 cells.

We next evaluated whether the computationally prioritized candidates exhibited anticancer activity in vitro. Selected candidates, excluding 15 previously reported compounds, were tested in the A375 and SK-MEL-5 melanoma cells, with dabrafenib as a reference drug. In SK-MEL-5 cells, YKL-06-062 and 4-PQBH markedly reduced cell viability to approximately 3.5% and 9.9%, respectively, while FL118-14-propanol, chimmitecan, CPX-351, and AZD-5597 also reduced cell viability below 50%. By contrast, all candidates tested in A375 cells yielded mean viabilities above 50% (Fig. 4b). Dose-response assays further showed that the IC₅₀ values of YKL-06-062 and 4-PQBH were 1.492 and 7.179 μM in SK-MEL-5 cells, compared with 27.41 and 41.94 μM in A375 cells (Fig. 4c). These measurements identified cell-line dependent responses among the prioritized candidates tested in the two melanoma models.

To further evaluate the generalizability of the model, we validated prioritized compounds in HepG2 and MCF7 cells. In HepG2 cells, HM43239 showed the strongest inhibitory effect, reducing cell viability to approximately 5%, followed by EM-651 (Fig. 4d). In MCF7 cells, most prioritized compounds showed limited activity, whereas DTCI exhibited a pronounced inhibitory effect (Fig. 4e). Dose-response assays confirmed the activity of HM43239 and EM-651 against HepG2 cells, with estimated IC₅₀ values of 0.642 μM and 4.29 μM, respectively (Fig. 4f). DTCI also showed concentration-dependent activity against MCF7 cells, with an estimated IC₅₀ of 0.902 μM, indicating greater potency than the reference drug doxorubicin (Fig. 4g). For compounds with high anticancer activity, we observed distinct pathway attention patterns across different drug candidates, suggesting cancer- and compound-specific mechanisms underlying their predicted activity. For example, in melanoma cells, YKL-06-062 prioritized several melanoma-relevant pathways, including melanogenesis, ErbB signaling, and ABC transporters. In contrast, HM43239 showed attention toward liver cancer-associated pathways, including VEGF signaling, Ras signaling, oxidative phosphorylation, and sphingolipid metabolism (Fig. S5).

In the reversal score-ranked list, 4-PQBH and YKL-06-062 were ranked 18th and 22nd, respectively (Table S4). Although the learned feature representations showed a tendency to separate effective and ineffective compounds, substantial overlap remained between the two groups, indicating that these features alone were insufficient for complete discrimination of compound efficacy (Fig. S6). We hypothesized that effective compounds would induce model-predicted perturbation profiles with greater separation from controls and therefore examined whether perturbation-control separation could provide additional information for identifying active compounds. The results showed that reported compounds and experimentally validated active candidates generally exhibited greater perturbation-control separation than inactive compounds, although some degree of overlap remained (Fig. S7a, Fig. S7c). Furthermore, both ARI and FMI values showed clear separation between active and inactive compounds, highlighting their potential as complementary metrics for prioritizing experimentally validated candidates (Fig. S7b). In line with this, 4-PQBH and YKL-06-062 achieved higher rankings when evaluated using these metrics, suggesting that incorporating perturbation-control separability can improve the identification of effective candidates.

### HM43239 exhibits marked antitumor activity against liver cancer

HM43239, also known as tuspetinib, is a multikinase inhibitor developed primarily for acute myeloid leukemia^31,32^. Our initial screening identified its inhibitory effect on HepG2 cell viability, prompting further evaluation in liver cancer cells and a mouse liver tumor model. To better characterize its inhibitory potency, we measured cell viability after 48 hours of treatment, using lenvatinib as a reference drug. HM43239 inhibited the viability of HepG2 and Huh7 cells more potently than lenvatinib, with IC_50_ values of 0.941 and 1.35 μM, respectively (Fig. 5a). Relative to untreated controls, HM43239 significantly reduced colony formation by 82.9% in Huh7 cells treated with 1 μM HM43239 and by 95.6% in HepG2 cells treated with 0.5 μM (Fig. 5b, Fig. S8a). Wound closure was also progressively reduced with increasing treatment concentrations. At 36 hours, wound closure was 79.35% in untreated cells, compared with 66.29%, 56.65%, and 41.32% in cells treated with 0.25, 0.5, and 1 μM, respectively (Fig. 5c, Fig. S8b). To assess whether apoptosis might contribute to the observed reduction in cell viability, we quantified apoptotic cells by Annexin V/PI staining. At the highest concentrations tested, the apoptotic population significantly increased from 8.36% to 57.8% in Huh7 cells and from 7.78% to 34.77% in HepG2 cells (Fig. 5d, Fig. S8c).

**Figure 5.**
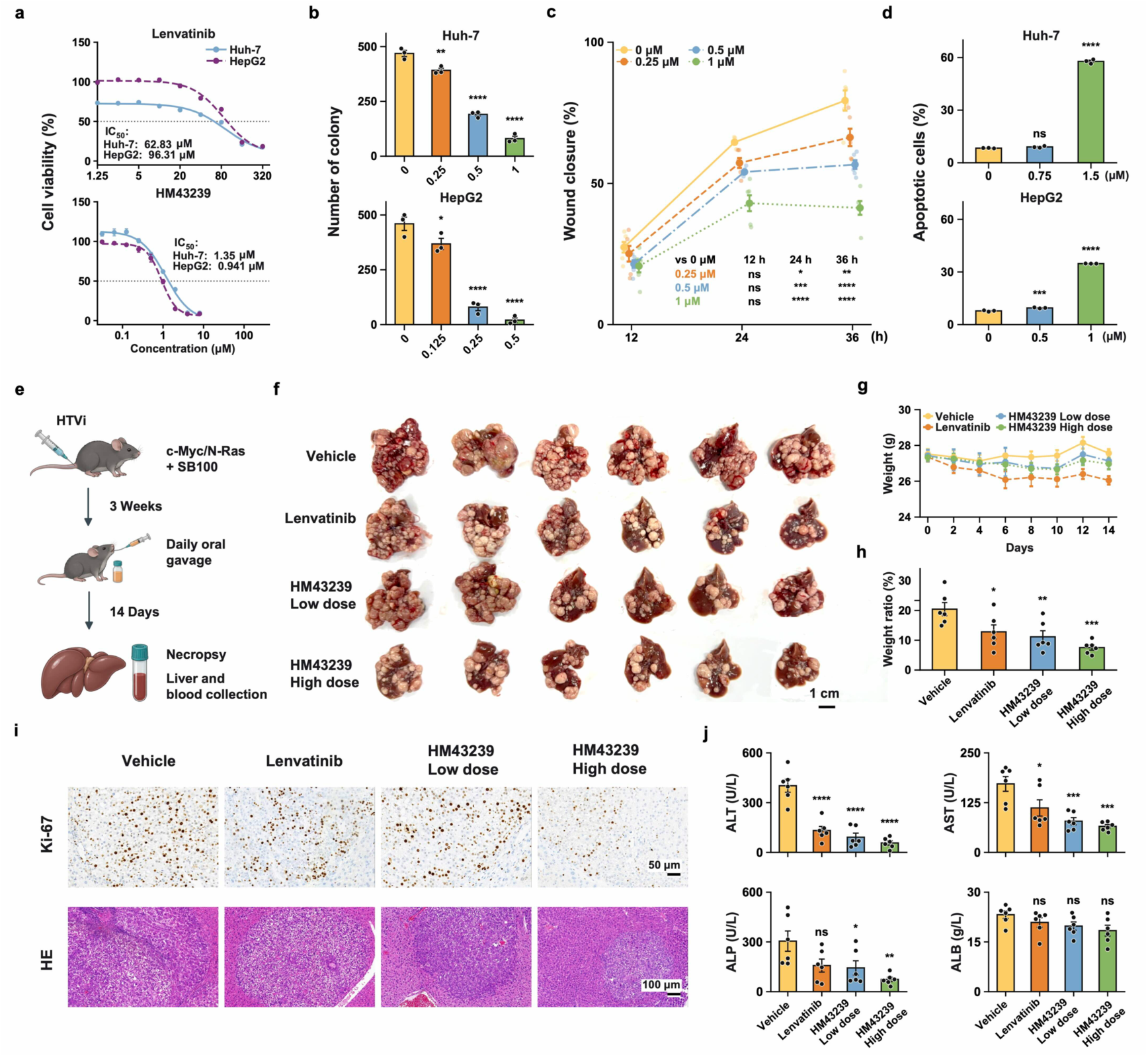
Evaluation of the efficacy and safety of HM43239. (a) Dose response curves for lenvatinib (top) and HM43239 (bottom) in Huh-7 and HepG2 cells. Blue solid and purple dashed curves represent Huh-7 and HepG2 cells, respectively. Dotted lines mark 50% cell viability, and the indicated IC₅₀ values were obtained from the fitted curves. (b) Colony counts of HM43239 treatment in Huh-7 and HepG2 cells. (c) Wound closure in Huh-7 cells at 12, 24, and 36 h after treatment with HM43239. (d) Percentages of apoptotic Huh-7 and HepG2 cells after HM43239 treatment, measured by Annexin V and PI staining. (e) Schematic of the animal experiments. Following hydrodynamic tail-vein delivery of c-Myc, N-Ras, and SB100 plasmids and three weeks of tumor induction, mice received vehicle, lenvatinib (4 mg/kg), low-dose HM43239 (5 mg/kg), or high-dose HM43239 (20 mg/kg) by daily oral gavage for 14 days. (f) Representative liver photographs at necropsy, with six mice shown per group. Scale bar, 1 cm. (g) Body weights during the 14-day treatment period. (h) Liver-to-body weight ratios at necropsy. (i) Representative liver sections stained for Ki-67 (top) or with H&E (bottom). (j) Serum levels of alanine aminotransferase, aspartate aminotransferase, alkaline phosphatase, and albumin at necropsy. Data comparisons were performed using one-way ANOVA followed by Dunnett’s test against the control group. *: p < 0.05, **: p < 0.01, ***: p < 0.001, ****: p < 0.0001.

Given the promising inhibitory effects of HM43239 observed in vitro, we next examined its antitumor activity in a mouse model of liver cancer driven by C-Myc and N-Ras (Fig. 5e). Following tumor induction, mice received daily oral treatment with vehicle, lenvatinib, low- or high-dose HM43239 for 14 days. Compared with vehicle-treated mice, those receiving HM43239 had fewer visible tumor nodules in the excised livers, particularly in the high-dose group (Fig. 5f). Body weight fluctuated modestly during treatment, with no marked decline, providing preliminary evidence of the safety of HM43239 at the tested doses (Fig. 5g). At the end of treatment, the mean liver-to-body weight ratios were significantly decreased, with 11.15% and 7.54% in the low- and high-dose HM43239 groups, respectively, compared with 20.43% in the vehicle group and 12.81% in the lenvatinib group (Fig. 5h). Ki-67 immunohistochemical staining was further performed to assess tumor cell proliferation, with representative sections showing fewer Ki-67-positive cells in the high-dose HM43239 group than in the vehicle group, suggesting reduced tumor cell proliferation following treatment (Fig. 5i).

We further evaluated serum biochemical indices and major-organ histology to characterize the effects of HM43239 treatment on liver function and tolerability. Serum ALT, AST, and ALP levels were lower in HM43239 treated mice than in vehicle group, consistent with an improvement in liver injury associated biochemical indices (Fig. 5j). Representative hematoxylin and eosin (H&E)-stained sections of the heart, spleen, lung, and kidney showed no obvious widespread structural abnormalities, and organ-to-body weight ratios were generally comparable across groups (Fig. S8d, Fig. S8e). Together with the observed antitumor activity, these findings provide support for the therapeutic potential of HM43239 in liver cancer.

### BioGCD discovers novel anti-melanoma compounds

To explore the potential of the model for novel drug discovery, we applied it to identify compounds without prior reports and selected the top 50 candidates for efficacy testing (Table S5). Z6412951658 was identified as an active hit, markedly inhibiting melanoma cell viability (Fig. 6a). Dose-response analysis further confirmed its anti-melanoma activity, yielding an IC_50_ value of 1.07 μM (Fig. 6b). Model-based interpretation using the KEGG pathway-prior attention weights revealed that Z6412951658 assigned high attention to pathways similar to those highlighted for YKL-06-062, such as melanogenesis and ErbB signaling pathways. This supports the pathway-level interpretability of our model and confirms that it can capture shared biological programs between functionally related compounds (Fig. S9a).

**Figure 6.**
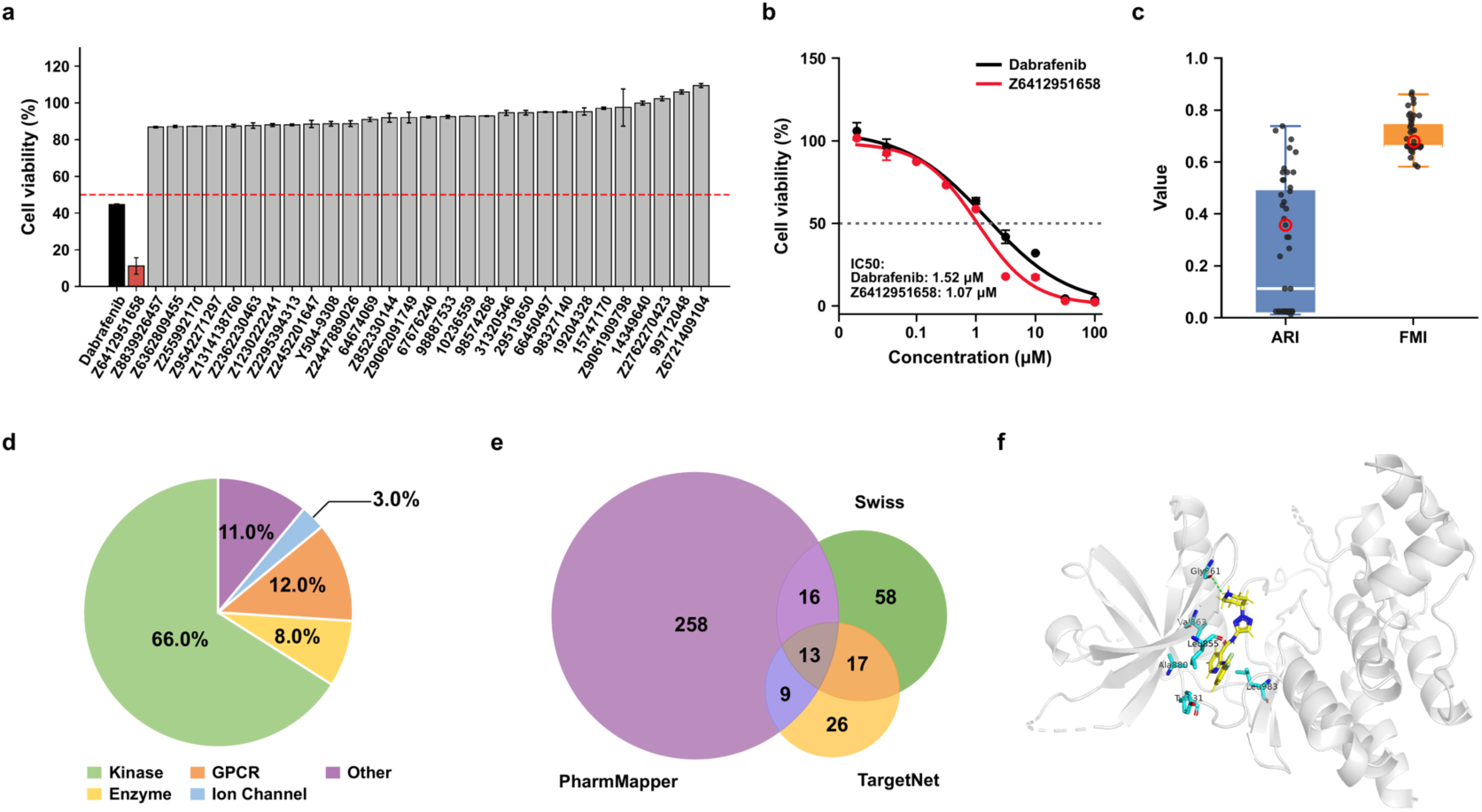
Model guided discovery of novel anti-melanoma candidates. (a) Cell viability screening of previously unreported candidates identified by the model. Dabrafenib serves as the reference compound, and Z6412951658 is highlighted in red as an active hit. (b) Dose response curves for Z6412951658 and dabrafenib in melanoma cells. (c) Distribution of ARI and FMI scores for model predicted perturbation-control separability across candidate compounds. Each point represents one compound, and red open circles identify Z6412951658. (d) SwissTargetPrediction classification of putative Z6412951658 targets. (e) Overlap among target sets predicted by PharmMapper, SwissTargetPrediction and TargetNet. (f) Predicted docking pose of Z6412951658 in the ATP-binding pocket of JAK2. The ligand is shown within the protein structure, with nearby residues and predicted contacts annotated.

Although marker gene reversal was used as the primary criterion for candidate prioritization, reversal strength alone was insufficient to clearly distinguish active from inactive compounds (Fig. S9b). We therefore assessed perturbation-control separability as a complementary criterion for candidate selection. Experimental validation showed that inactive compounds tended to have low values in at least one of the two perturbation-control separability metrics, ARI or FMI. In particular, compounds with extremely low ARI or FMI values were consistently inactive, further confirming that these metrics can serve as complementary filters to deprioritize low-confidence candidates and reduce the number of compounds requiring experimental testing during novel drug discovery (Fig. 6c, Table S5).

Target prediction of Z6412951658 was performed using multiple computational approaches. SwissTargetPrediction identified a broad range of candidate targets, with the majority belonging to kinase-related proteins (66%), followed by GPCRs (12%), enzymes (8%), ion channels (3%), and other target classes (11%) (Fig. 6d, Table S6). The overlap analysis among SwissTargetPrediction, TargetNet, and PharmMapper revealed 13 shared candidate targets, among which JAK2 was identified as a melanoma-associated target involved in tumor progression and therapeutic response^33,34^ (Fig. 6e, Table S6). Molecular docking analysis revealed a potential binding mode of Z6412951658 with JAK2. Z6412951658 was positioned within the ATP-binding pocket of the JAK2 kinase domain, forming a hydrogen bond with Gly861 and hydrophobic interactions with surrounding residues, including Leu855, Val863, Ala880, Tyr931, and Leu983 (Fig. 6f). The predicted binding mode showed that Z6412951658 could stably occupy the ATP-binding cavity and establish hydrogen bonding and hydrophobic interactions with key residues, supporting JAK2 as a potential binding target underlying its anti-melanoma activity.

### BioGCD enables prediction of synergistic compound combinations

We further evaluated whether BioGCD predictions could be used to inform the selection of two compound combinations. Compound-pair gene coverage and correlations between compound induced transcriptional responses have been shown to inform compound combination prediction^35,36^. We therefore analyzed three melanoma cell lines and compared synergistic and non-synergistic compound pairs using three features: the overlap ratio of the top 10 perturbed genes, the Pearson correlation of compound induced gene-expression changes and the Jensen-Shannon similarity of pathway-level responses, an output of the BioGCD pathway module that captures pathway-level differences between compound pairs. These features showed cell-line dependent differences between synergistic and non-synergistic pairs, with no consistent direction of change across the three melanoma cell lines (Fig. 7a). Similar heterogeneity was observed across breast cancer cell lines (Fig. S10), suggesting that genetic and transcriptional differences among cell lines preclude the use of any single feature as a universal indicator of compound combination efficacy.

**Figure 7.**
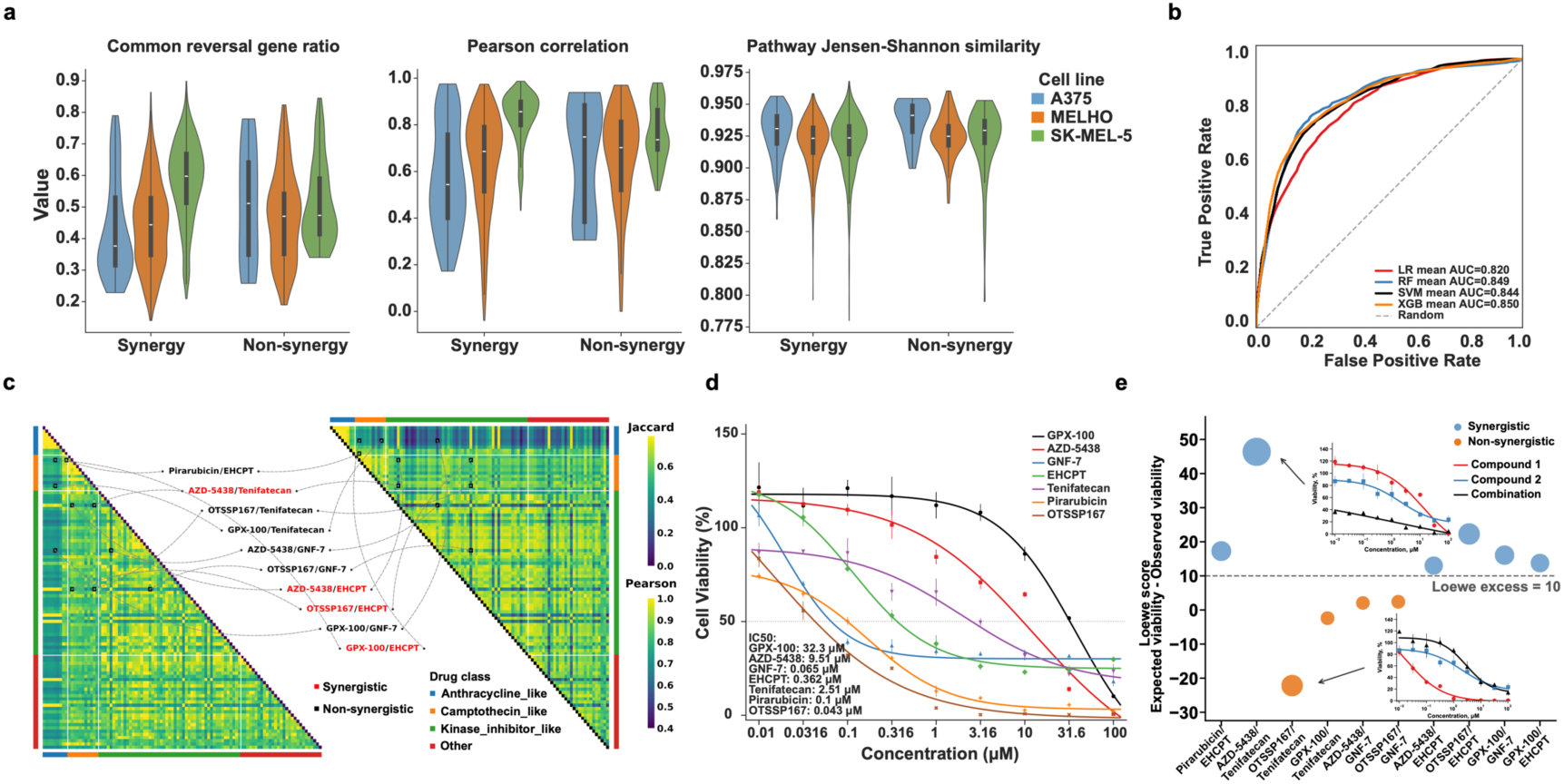
Identification and validation of synergistic compound combinations in melanoma cells. (a) Distributions of the common reversal-gene ratio, Pearson correlation between predicted transcriptional responses, and pathway Jensen-Shannon similarity across synergistic and non-synergistic compound pairs. (b) ROC analysis for assessing the ability of computational features to distinguish synergistic from non-synergistic combinations. (c) Heatmaps of the reversal-gene Jaccard index and Pearson correlation among candidate compounds. Color strips annotate compound classes, and connecting lines denote the ten compound pairs selected for experimental testing. Red labels indicate pairs predicted to be synergistic, whereas black labels indicate pairs predicted to be non-synergistic. (d) Dose response curves and IC₅₀ values of seven individual candidate compounds. (e) Experimental Loewe synergy scores for the ten selected compound pairs in SK-MEL-5 cells. Blue and orange points indicate experimentally synergistic and non-synergistic combinations, respectively, using the dashed threshold of 10. Insets show dose response curves of representative compounds.

Motivated by these observations, we expanded aforementioned measurements into a broader feature set and constructed classification models for melanoma compound-combination prediction using either pooled melanoma cell lines or SK-MEL-5 alone. Consistent with the cell-line dependent feature distributions, the model trained on pooled melanoma cell lines showed limited predictive performance for two-compound combinations, with all classifiers yielding average AUC values above 0.70 and XGBoost achieving the highest AUC of 0.75, probably reflecting confounding by inter-cell-line heterogeneity (Fig. S11a). By contrast, the SK-MEL-5 specific model achieved substantially improved classification performance, with all AUC values exceeding 0.80 (Fig. 7b and Fig. S11b).

To experimentally evaluate nominated combinations, we selected ten predicted compound pairs, including six predicted negative and four predicted positive combinations, and performed single-agent and combination-response assays in SK-MEL-5 cells (Fig. 7c). Among the seven single agents tested, GPX-100, AZD-5438 and Tenifatecan exhibited relatively weak single-agent activity (Fig. 7d). According to Loewe synergy scores, all four combinations predicted as positive were experimentally validated as synergistic, but two of the six combinations predicted as negative (Pirarubicin/EHCPT and GPX-100/GNF-7) also showed synergy in the validation assays (Fig. 7e). Notably, the approach was able to identify combinations that were not evident from single-agent activity alone, including AZD-5438/Tenifatecan, which was correctly predicted and experimentally confirmed as synergistic.

## Discussion

In this study, we developed BioGCD, a biologically grounded multimodal deep learning framework for predicting compound induced transcriptional responses and prioritizing therapeutic candidates. By integrating gene-level association graphs, chromosomal proximity, GO annotations, KEGG pathways, transcription factor-target interactions, and a compound-aware pathway attention encoder, this approach incorporates biological prior knowledge at both gene and pathway levels. Under compound-disjoint benchmarking, the model consistently outperformed representative existing methods, demonstrating robust generalization to unseen compounds. In addition, it can predict transcriptional responses across different compound concentrations and more accurately recovered biologically meaningful perturbation signatures, supporting its ability to capture both dose-dependent and mechanism-relevant perturbational effects.

The framework showed substantial value for drug repurposing across multiple cancer contexts. In melanoma and hepatocellular carcinoma cells, 21 of the top 30 prioritized compounds were validated experimentally or supported by literature evidence, while in breast cancer cells, 9 of the top 30 candidates showed inhibitory activity. Among these candidates, HM43239 is a multi-kinase inhibitor targeting FLT3 and other leukemia-relevant kinases that was developed primarily for acute myeloid leukemia. In this study, HM43239 exhibited potent activity against liver cancer cells, reducing cell viability, colony formation, and wound closure while promoting apoptosis. It also reduced tumor burden in a mouse liver tumor model, with no obvious widespread structural abnormalities in representative sections of the examined major organs after treatment, highlighting its potential for liver cancer treatment and the ability of our model to uncover cross-indication drug repurposing opportunities.

Beyond repurposing, BioGCD enabled de novo drug discovery. Among the top 50 candidates without prior biological annotation, Z6412951658 was experimentally confirmed as a potent anti-melanoma hit, inhibiting cell viability with an IC₅₀ of 1.07 μM and exhibiting greater potency than the positive control. Integrated target prediction and molecular docking suggested JAK2 as a potential target, with stable hydrogen-bond and hydrophobic interactions within the ATP-binding pocket. This finding provides both a novel chemical starting point and a mechanistic hypothesis for subsequent validation and medicinal chemistry optimization.

We further extended BioGCD to recommend synergistic compound combinations. Features derived from compound-perturbation predictions indicated that combination signatures are highly context dependent, as transcriptional and pathway-level features distinguished synergistic from non-synergistic pairs in a cell-line-specific manner but did not show consistent trends across melanoma or breast cancer models. This heterogeneity suggests that single response-derived metrics are unlikely to serve as universal predictors of synergy. Instead, predictions-derived features were more informative when incorporated into cell-line-specific models, as illustrated by the improved performance of the SK-MEL-5 model and its ability to nominate experimentally validated synergistic pairs, including AZD-5438/Tenifatecan. This highlights the importance of incorporating cellular context into BioGCD-based combination prediction.

The model is not simply a black-box predictor. Attribution analyses revealed structure-dependent molecular patterns consistent with known pharmacophoric features, as illustrated by vemurafenib, linking specific chemical substructures to downstream transcriptional responses. Meanwhile, pathway attention distinguished mechanism-relevant pathways from unrelated ones and demonstrated that the learned pathway representations retained biological interpretability. This structure- and pathway-level interpretability helps move BioGCD beyond black-box prediction toward more mechanism-aware modeling.

Although BioGCD demonstrated better predictive performance and biological interpretability, several limitations should be considered. First, the model was primarily trained on standardized L1000 transcriptomic profiles, which may limit its generalizability to other experimental platforms, such as single-cell sequencing. Second, while attribution analysis and pathway-attention mechanisms enhanced model interpretability, the resulting insights remain computational inferences that require causal validation through targeted genetic or pharmacological perturbations. Future work will incorporate time-series modeling and additional omics modalities, including proteomics and epigenomics, to further improve the biological resolution of the model. In addition, its performance should be evaluated in models that more closely recapitulate human physiological conditions, such as patient-derived organoids and in vivo tumor xenografts. These efforts may broaden the applicability of BioGCD across disease contexts, facilitate systematic validation of predicted synergistic compound combinations, and support the development of personalized combination therapy strategies.

## Methods

### L1000 data curation and preprocessing

Gene-expression profiles were obtained from the L1000 perturbation dataset. Samples generated after 24 h of compound exposure were retained. Replicate profiles corresponding to the same cell line and compound were averaged during upstream data curation to generate a single perturbation profile for each condition.

Profiles failing the predefined upstream quality-control criteria were removed. The curated dataset contained 100,596 perturbation profiles and 164 vehicle-control profiles representing 164 cell lines, with expression measurements for 12,328 genes per profile. During model specific preprocessing, perturbation and control matrices were aligned using their shared gene identifiers and reordered to ensure an identical gene order. For each perturbation profile, expression variance was calculated across the aligned gene set, and profiles with a variance of 1 × 10^-^⁴ or lower were excluded.

The model training pipeline used the expression scale stored in the processed input files without applying additional library-size normalization or log transformation. Before tensor construction, non-finite values, including NaN and positive or negative infinity, were replaced with zero, and negative expression values were clipped to zero.

### Compound-disjoint data partitioning and control matching

To evaluate prediction for previously unseen compounds, the dataset was partitioned at the compound level rather than the sample level. Unique compounds were defined by their canonical SMILES representations, randomly shuffled, and divided into training, validation, and test sets at an 8:1:1 ratio. Consequently, all perturbation samples associated with the same compound were assigned to the same subset. The entire data-splitting and model-training procedure was independently repeated three times using different random seeds to assess the robustness of model performance.

For each perturbation profile, a vehicle-control profile from the same cell line was selected. Samples without an available cell-line-matched control were excluded. The training, validation, and test pairing tables were stored to preserve the exact perturbation-control assignments.

### Molecular and cellular representations

Compounds were represented using 1,024-bit functional-class circular fingerprints with a diameter of four (FCFP4). SMILES strings were parsed using RDKit and molecular fingerprints were generated using a Morgan radius of 2 with feature-based atom invariants.

Cellular context was represented by the unperturbed gene-expression profile together with a precomputed 1,280-dimensional Universal Cell Embedding (UCE) obtained using a pretrained foundation model. The UCE representation was stored with each control profile and used as a fixed input feature without further fine-tuning of the UCE model.

Compound concentration was encoded from a scalar value into the 1,024-dimensional molecular feature space using a two-layer dose encoder (1→64→1,024, with a ReLU activation). The resulting dose embedding was added element-wise to the molecular fingerprint before tokenization and attention-based fusion with the cellular representation.

### Biological graph-prior encoder

We constructed four gene-level prior graphs encoding chromosome-positional relationships, Gene Ontology (GO)-based functional relationships, KEGG-derived gene relationships, and transcription factor-target gene (TF-TG) regulatory interactions. The chromosome-positional prior followed the strategy described in Cell-GraphCompass^37^. The GO prior was constructed from Gene Ontology annotations following the GEARS-based strategy^38,39^. KEGG relationships were derived from KEGG Release 119.0^26^ and TF-TG regulatory relationships were obtained from the TFTG^40^ and TRRUST^41^ databases.

For each graph r, edges below the graph-specific importance threshold were removed (0.1 for the KEGG graph and 0 for the other three graphs), and up to the 21 highest-weight incoming edges were retained per target gene. During adjacency-matrix construction, edges whose source or target genes were absent from the aligned 12,328 gene expression space were discarded. A self-loop with weight 1 was added to each participating gene lacking one. The adjacency matrix used target genes as rows and source genes as columns and was row-normalized by the total incoming edge weight of each target gene.

Let E denote the learnable gene embedding matrix of size G × H, where G = 12,328 and H = 1,024. For each prior graph r, a single graph-propagation step aggregated source gene embedding into their target genes using the normalized adjacency matrix, followed by a graph-specific transformation:

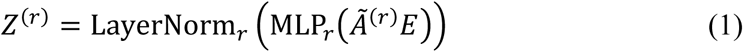

where each graph-specific MLP comprised a linear layer, ReLU activation, dropout (rate 0.1), and a second linear layer, followed by graph-specific layer normalization. The four graph representations were combined using learnable scalar sigmoid gates:

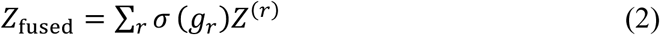

where each scalar gate logit was initialized to −2.0. The fused representation was then subjected to layer normalization and dropout (rate 0.1):

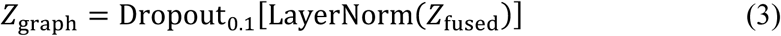

To derive a sample-specific graph context, baseline expression values were clipped to non-negative values and normalized by the total non-negative expression of each sample. For numerical stability, the denominator was bounded below by 10⁻⁶:

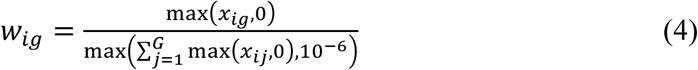

The sample-specific graph representation was then calculated as

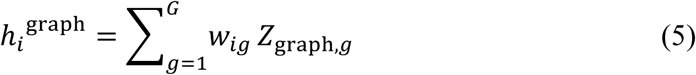

If a sample had zero total non-negative expression, all expression weights and the resulting graph context were zero. The H-dimensional graph-context vector was then concatenated with the baseline expression profile and the 1,280-dimensional UCE representation before input projection.

### KEGG pathway-prior module

A separate pathway-prior module incorporated gene-to-pathway membership information derived from KEGG. Two sparse membership matrices were constructed. The gene-to-pathway matrix, **M**_a→c_ ∈ ℝ*^P^*^×*G*^, aggregated gene expression into pathway activities by assigning each member gene a weight equal to the reciprocal of the number of genes in the corresponding pathway. Conversely, the pathway-to-gene matrix, **M**_c→a_ ∈ ℝ*^G^*^×*P*^, mapped pathway-level changes back to genes by averaging across the pathways associated with each gene.

For sample *i* with baseline expression vector **x***_i_*, pathway activity was calculated as

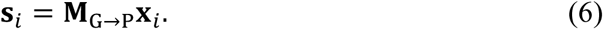

Each scalar pathway activity *s_ip_* was projected into a 128-dimensional representation using a linear layer followed by ReLU activation and layer normalization, and was then combined with a learnable pathway-specific embedding. Dropout was subsequently applied to obtain the pathway token **t***_ip_*:

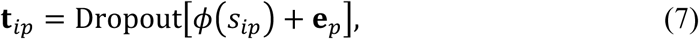

where *φ*(⋅) denotes the activity projection and **e***_p_* is the learnable embedding of pathway *p*.

The dose-adjusted molecular representation was linearly projected into the same 128-dimensional space to form a compound query **q***_i_*. Compound-aware pathway attention weights were then calculated using scaled dot-product attention:

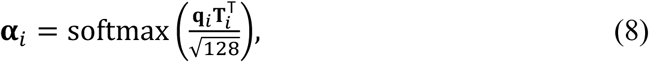

where **T***_i_* contains the pathway tokens of sample *i*.

A pathway-specific expression-change value was independently estimated from each pathway token using a two-layer multilayer perceptron and modulated by its corresponding compound-aware attention weight:

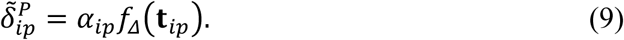

The resulting pathway-level changes were mapped back to the gene space using the pathway-to-gene matrix:

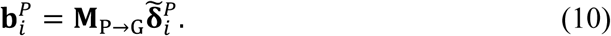

Finally, the pathway-derived gene-level bias was incorporated into the expression-change prediction from the main decoder through a learnable sigmoid gate:

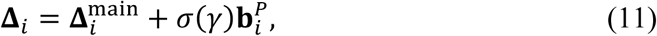

where the gate logit *γ* was initialized to −4.0. The final perturbed expression profile was obtained using a residual formulation,

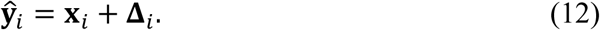

### BioGCD architecture

BioGCD received four inputs for each perturbation condition: the baseline gene-expression profile, the UCE cell embedding, the molecular fingerprint, and the compound concentration. The baseline expression vector, UCE embedding, and sample-specific graph-prior representation were concatenated and projected into a 1,024-dimensional cellular representation. In parallel, the scalar compound concentration was transformed by the dose encoder and added to the 1,024-dimensional molecular fingerprint, yielding a dose-adjusted molecular representation of the same dimensionality.

Both the cellular and molecular representations were reshaped into eight tokens of 128 dimensions. Cross-attention was then applied using the cellular tokens as queries and the molecular tokens as keys and values:

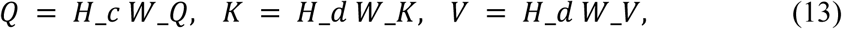

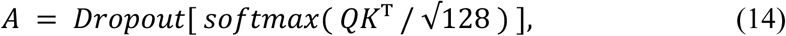

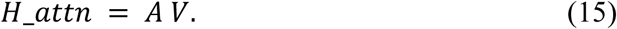

The attention output was flattened into a 1,024-dimensional vector and processed by a fusion multilayer perceptron consisting of a linear layer, ReLU activation, layer normalization, dropout, and a second linear layer. The resulting fused representation was passed to a decoder with hidden dimensions of 2,048 and 1,024, followed by an output layer matching the aligned gene-expression dimension (G = 12,328). The decoder therefore produced a gene-level compound induced expression-change vector, Δ_decoder. A dropout rate of 0.1 was used in the attention, fusion, and decoder components, with ReLU activations applied to the corresponding hidden layers.

The KEGG pathway-prior module provided an additional gene-level expression-change bias, which was incorporated into the decoder output through a learnable sigmoid gate before residual reconstruction. The final perturbed expression profile was calculated as

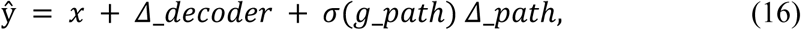

where x denotes the baseline control-expression profile and g_path is the learnable pathway-gate logit.

An auxiliary classification head was applied to the fused representation to predict cell-type identity. The classification head consisted of a 256-unit hidden layer with ReLU activation and dropout, followed by an output layer corresponding to the number of cell types.

### Model training and selection

BioGCD was implemented in PyTorch. Models were trained for 200 epochs using the Adam optimizer with a learning rate of 1 × 10^−4^, a batch size of 256, and gradient clipping with a maximum norm of 5.0.

The expression-prediction objective was defined on compound induced expression changes rather than on absolute expression levels:

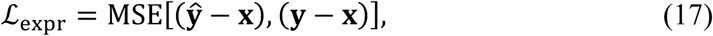

where **x** denotes the baseline control-expression profile, **y** the observed perturbed expression profile, and **ŷ** the model prediction.

The total training objective combined the expression loss with an auxiliary cell-type classification loss:

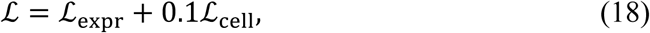

where ℒ_cell_ was the cross-entropy loss for cell-type classification.

Model selection was based on the mean sample-wise Pearson correlation coefficient on the validation set. Pearson correlation was calculated across genes for each validation sample and then averaged across all valid samples. The checkpoint achieving the highest mean validation Pearson correlation was retained. After training, this checkpoint was reloaded and evaluated once on the held-out test set.

### Evaluation metrics

Predictive performance was quantified for each held-out perturbation profile over all G = 12,328 genes. Let y_i and ŷ_i denote the observed and predicted post-treatment expression vectors for sample i, respectively, and let x_i denote its matched control vector. Pearson correlation coefficient (PCC), Spearman rank correlation coefficient (SCC), coefficient of determination (R²) and mean squared error (MSE) were first computed from the absolute post-treatment expression profiles. PCC was calculated as

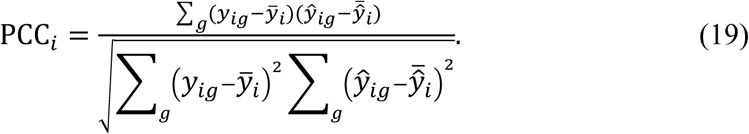

SCC was calculated as the PCC between the within-sample ranks of the observed and predicted gene values:

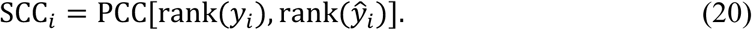

R² and MSE were calculated for each sample as

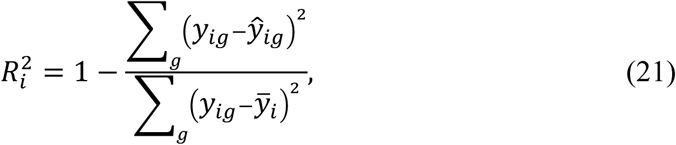

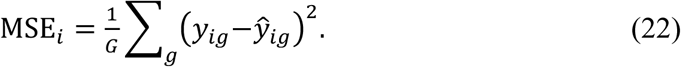

To evaluate compounds induced responses independently of baseline expression, the observed and predicted expression-change vectors were defined relative to the same matched control:

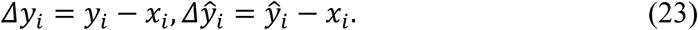

The four change-based metrics, denoted ΔPCC, ΔSCC, ΔR² and ΔMSE, were obtained by applying the definitions above to Δy_i and Δŷ_i. Higher PCC, SCC and R² values indicate better agreement, whereas lower MSE values indicate smaller prediction errors. Non-finite PCC, SCC or R² values arising from zero-variance profiles were assigned a value of zero, consistent with the evaluation implementation.

Performance was also evaluated on the 100 genes showing the largest observed perturbation response. For each cell-line compound condition, expression values were averaged across replicate samples, and observed log₂ fold changes were calculated relative to the corresponding mean control profile:

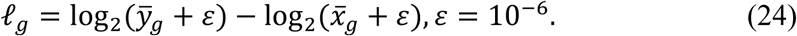

Genes were eligible for selection when max(y̅_g, x̅_g) ≥ 0.1. Among eligible genes, the 100 genes with the largest |ℓ_g| were defined as D_100:

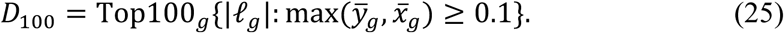

The same condition-specific D_100 set was applied to every sample from that condition. PCC, SCC, R² and MSE, together with their four change-based counterparts, were then recomputed after restricting y_i, ŷ_i, Δy_i and Δŷ_i to D_100.

Directional recovery was evaluated using the top 100 upregulated and top 100 downregulated genes derived separately from the observed and predicted log₂ fold-change vectors. If T_i^up and P_i^up denote the observed and predicted upregulated sets, and T_i^down and P_i^down the corresponding downregulated sets, the directional overlap percentages were

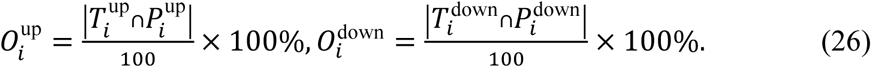

For the representative perturbation conditions shown in the log₂ fold-change scatter plots, PCC and SCC were additionally calculated between observed and predicted log₂ fold changes over D_100. Genome-wide direction accuracy was calculated as the percentage of genes for which the predicted and observed expression changes had the same sign:

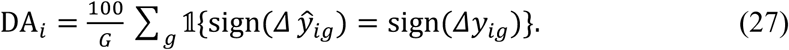

The expression-based metrics were calculated at the sample level and averaged within each held-out test split. Bar plots report the mean and standard deviation across three independently generated compound-disjoint splits. Direction-accuracy distributions were summarized across cell-line compound conditions pooled from the three splits. For radar visualization only, the model-level mean for each metric was min-max scaled across the compared models; MSE axes were reversed so that larger normalized values consistently represented better performance:

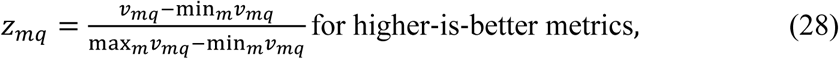

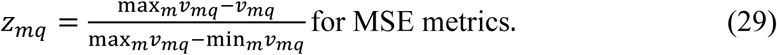

Here, v_mq denotes the mean value of metric q for model m. If all compared models had the same value for a metric, the normalized value was set to 1 for every model.

### Transcriptome-wide Assessment

Transcriptome-wide impact and the effective number of differentially expressed genes were calculated according to the TRADE framework^42^. Transcriptome-wide impact, TI, was defined as the variance of the true differential expression effects across all measured genes:

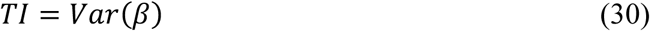

where *β* denotes the gene-level differential expression effect. A larger *TI* indicates a stronger global transcriptional effect of a perturbation.

The effective number of differentially expressed genes, πDEG, was defined as:

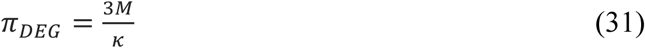

where *M* is the number of measured genes and *κ* is the kurtosis of the true effect distribution:

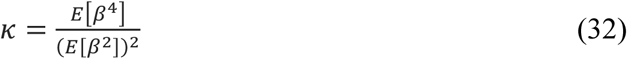

*πDEG* reflects the breadth of transcriptional effects across the transcriptome.

### Cancer-associated transcriptional signatures

For hepatocellular carcinoma and breast cancer, disease signatures were derived from TCGA tumor-normal comparisons. Genes with an FDR of <0.05 and an absolute log₂ fold change of ≥3 were retained. For melanoma, a treatment-response signature was aggregated from three EMBL-EBI studies of BRAF inhibitor-treated melanoma cells and their corresponding controls, retaining genes with an absolute log₂ fold change of >3. After removing duplicate and directionally conflicting genes, all signatures were intersected with the 12,328-gene BioGCD feature space. Because the source datasets differed in experimental design, these signatures were used only as cancer-specific references for within-cancer compound prioritization and were not compared quantitatively across cancer types.

### Reversal score calculation

To quantify the extent to which a predicted compound induced transcriptional response reversed the disease-associated expression signature, disease-associated upregulated and downregulated gene sets were first processed by removing genes present in both sets. The remaining genes were restricted to those represented in the model output. For each candidate compound *d*, all measured genes were ranked in descending order of predicted log_2_FC, such that rank 1 corresponded to the largest predicted log_2_FC value. Ties were assigned according to their order in the expression matrix.

For a query gene set *Q* containing *p* genes, let *n* denote the total number of genes in the predicted expression profile and let *r*_(*m*),*d*_ denote the *m*-th smallest rank among the genes in *Q* for compound *d*. Two rank-based enrichment components were calculated as

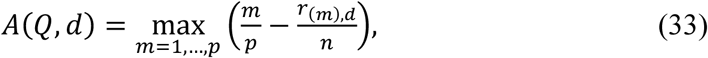

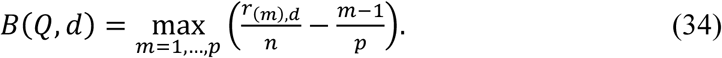

The enrichment score for query set *Q* was defined directly as

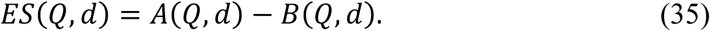

Separate enrichment scores were calculated for the disease-upregulated and disease-downregulated gene sets, denoted *Q*_up_ and *Q*_down_, respectively. The final reversal score for compound *d* was calculated as

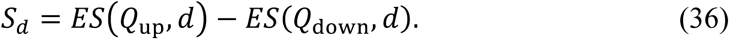

Because genes were ranked from high to low predicted log_2_FC, a strong reversal placed disease-upregulated genes toward the lower end of the compound induced ranking and disease-downregulated genes toward the upper end. Consequently, lower values of *S_d_* indicated stronger predicted reversal of the disease-associated transcriptional signature, and candidate compounds were ranked in ascending order of *S_d_*. Scoring was performed only when at least five genes from each of the filtered upregulated and downregulated query sets were represented in the model output.

### Compound libraries

Two compound libraries supplied by TargetMol (TaoSu Biotech) were used. The T001 library, comprising 20,674 compounds, served as a focused collection for compound prioritization and subsequent experimental validation. To expand the chemical search space, we additionally used a large-scale screening collection comprising 6,351,767 compounds curated from the TargetMol database. This collection was divided into “core compounds”, including molecules in ongoing clinical trials or discontinued after failing to meet clinical endpoints, and “selected compounds”, for which no prior biological annotation was available in the source records.

### Single compound prioritization

Single compound prioritization was performed for each target cell line using the TargetMol compound library. Compounds with missing or invalid SMILES representations were excluded before prediction.

For each valid compound, BioGCD predicted the post-treatment gene expression profile at 10 μM. When multiple control profiles were available for the target cell line, predictions were generated independently for each control. For gene *g*, compound *d*, and control profile *c*, the predicted log2 fold change was calculated as

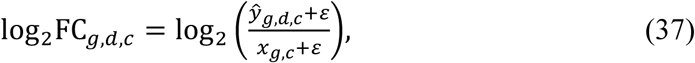

where *ε* = 10^−6^, *ŷ_g,d,c_* denotes the predicted perturbed expression, and *x*_g,*c*_ denotes the corresponding baseline expression. When multiple controls were available, log2FC values were calculated separately relative to each control and then averaged:

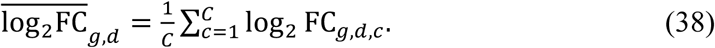

Disease-associated upregulated and downregulated gene sets were used as the query signature. Genes occurring in both sets were removed, and the remaining genes were restricted to those represented in the model output. For each compound, genes were ranked in descending order of predicted log2FC.

Reversal scores were calculated using the rank-based enrichment procedure defined in Eqs. (33)-(36).

Lower reversal scores indicated stronger predicted reversal of the disease-associated transcriptional signature, corresponding to disease-upregulated genes being shifted toward compound-induced downregulation and disease-downregulated genes toward compound induced upregulation. Compounds were ranked in ascending order of *S_d_*. The top 30 ranked candidates were subsequently selected for downstream analysis and experimental validation.

### Compound synergy prediction

For each target cell line, we first selected the top 100 compounds ranked by the predicted reversal score from the single-compound prioritization results. Candidate two-compound combinations were then evaluated using features designed to capture gene-level complementarity and transcriptional response relationships between compound pairs. These features included the Jaccard index of the top 10 percent perturbed genes for the two compounds, the Pearson correlation of marker genes, fold-change-based relationships, pathway-level similarity, and molecular Tanimoto similarity.

To train the compound-combination synergy prediction model, we used combination response data from DrugComb^43^. Compounds that were included in the L1000 training set were excluded to avoid overlap with compounds used for model training. Among the remaining compound pairs, combinations with a Loewe synergy score greater than or equal to 10 were labeled as synergistic positive examples, whereas those with a Loewe synergy score below 10 were labeled as negative examples. The labeled data were split into training and validation sets at a 4:1 ratio. To assess robustness, data splitting was repeated with 20 random seeds, producing 20 independent training and validation sets.

For each dataset, we trained four classification models, including logistic regression, random forest, support vector machine, and extreme gradient boosting. Ensemble prediction was performed across the 20 independently trained models. A compound pair was classified as synergistic only if it was predicted to be synergistic by at least 10 models.

### Cell culture

All cell lines used in this study were obtained from the China Center for Type Culture Collection. A375, HepG2, Huh-7, MCF7, and SK-MEL-5 cells were cultured in 10 cm culture dishes containing 10 mL of complete Dulbecco’s modified Eagle medium (DMEM) supplemented with 10% fetal bovine serum and 1% penicillin-streptomycin. Cells were maintained at 37.5 °C in a humidified incubator with 5% CO₂. When cells reached approximately 70%-80% confluence, they were detached with 0.25% trypsin-EDTA and passaged.

### Cell viability assays

Cells were seeded into 96-well plates at 5 × 10³ cells/well in 100 μL of complete medium and incubated at 37.5 °C with 5% CO₂. Compounds were prepared as 10 mM DMSO stocks and diluted in complete medium to the indicated concentrations. Vehicle-control wells received the same final concentration of DMSO. After adding 150 μL of compound-containing medium to each well, cells were treated for 24 h or 48 h. After treatment, the medium was replaced with 100 μL of complete medium containing 10% (v/v) CCK-8 reagent (TargetMol, C0005). Plates were incubated in the dark for 1 h at 37.5 °C with 5% CO₂, and absorbance was measured at 450 nm.

### Colony Formation Assay

Huh-7 and HepG2 cells were seeded in six-well plates at a density of 5 × 10³ cells per well. Huh-7 cells were treated with HM43239 at concentrations of 0, 0.25, 0.5, or 1 μM, whereas HepG2 cells were treated with 0, 0.125, 0.25, or 0.5 μM. After 14 days of culture, the colonies were fixed with 4% paraformaldehyde and stained with 0.5% crystal violet for 30 min. Excess stain was removed with distilled water, and the plates were allowed to dry. The colonies were then photographed and quantified.

### Wound-Healing Assay

Huh-7 cells were seeded in six-well plates and cultured to 90%-95% confluence. A scratch was made with a sterile 200 μL pipette tip, and cells were washed three times with PBS to remove detached cells. Cells were then treated with 0, 0.25, 0.5, or 1 μM HM43239. Images were captured using an inverted microscope at 0, 12, 24, and 36 h after treatment. Wound areas were measured using ImageJ, and cell migration was assessed based on the change in wound area.

### Animal Experiments

Six-week-old C57BL/6JGpt mice were purchased from GemPharmatech Co., Ltd. To establish the HCC model, c-Myc, N-Ras, and SB100 transposase plasmids were mixed in 2 mL of sterile 0.9% saline and injected via the tail vein within 6-7 s. Three weeks later, mice were randomly assigned to vehicle, HM43239 (5 or 20 mg/kg), or lenvatinib (4 mg/kg) groups, with six mice per group. Treatments were administered daily by oral gavage, and body weight was measured every two days throughout treatment. At the predefined endpoint, mice were euthanized, and the liver and other major organs were collected. Liver weight was measured to calculate the liver-to-body weight ratio. All animal experiments were approved by the relevant institutional animal ethics committee (Approval number: CPUTML-HWW-260803-01).

### H&E Staining and Immunohistochemistry

Tissues were fixed in 4% paraformaldehyde, dehydrated, embedded in paraffin, and sectioned. H&E staining was performed using a standard automated procedure. For immunohistochemistry, sections were deparaffinized, rehydrated, and subjected to heat-induced antigen retrieval in citrate buffer. After cooling, sections were incubated with an anti-Ki-67 primary antibody overnight at 4 °C, washed, and incubated with an HRP-labeled polymer anti-rabbit secondary antibody for 20 min at room temperature. Staining was visualized using 3,3′-diaminobenzidine (DAB).

### Serum Biochemical Analysis

Serum samples were collected at the experimental endpoint. Biochemical parameters related to liver and kidney function, metabolism, and tissue injury were measured using routine clinical laboratory methods. The in vivo tolerability of HM43239 was assessed in conjunction with organ indices and histopathological findings.

### Statistical analysis

Data were analyzed using one-way ANOVA or mixed two-way repeated-measures ANOVA, followed by appropriate multiple-comparison tests, as specified in the figure legends. *p* < 0.05 was considered statistically significant.

### Experimental validation of compound synergy

Compound combinations were tested using a fixed-ratio design based on the single-agent IC₅₀ values. For each combination, the IC₅₀ ratio of Compound A to Compound B was used to define the combination concentrations. Compound A was tested at nine serial concentrations: 100, 31.6, 10, 3.16, 1, 0.316, 0.1, 0.0316, and 0.01 μM, while Compound B was diluted in parallel according to the fixed IC₅₀ ratio. Each concentration was tested in triplicate, with blank and negative-control wells included. Cell viability was measured as described above. The IC₅₀ of each fixed-ratio combination was estimated from the combination dose-response curve, and the fixed-ratio Loewe synergy score was calculated according to the Loewe additivity model.

## Data availability

All datasets analyzed in this study are publicly available. L1000 expression profiles were obtained from the Gene Expression Omnibus (GEO) under accession number GSE92742. RNA-seq data from The Cancer Genome Atlas (TCGA) were downloaded from the Genomic Data Commons (GDC) Data Portal (https://portal.gdc.cancer.gov/). Compound-combination response data were obtained from the DrugComb database (https://drugcomb.org/).

## Code availability

The code for BioGCD is available at https://github.com/wenjean/BioGCD.

## Author contributions

J.W. contributed to conceptualization, methodology, investigation, software, formal analysis, validation, data curation, writing the original draft, and visualization. Z.X. contributed to data curation, formal analysis, investigation, supervision, validation and writing the original draft. S.Y., X.Z., C.W. and K.G. contributed to experimental validation and analysis of experimental results. Z.L., W.H., J.W, L.H., W.P. and Y.D. contributed to investigation and formal analysis. A.L., H.W. and Y.Y. contributed to conceptualization and project administration. C.P. contributed to conceptualization, methodology, project administration, funding acquisition, and review and editing of the manuscript.

---

**Figure S1.**
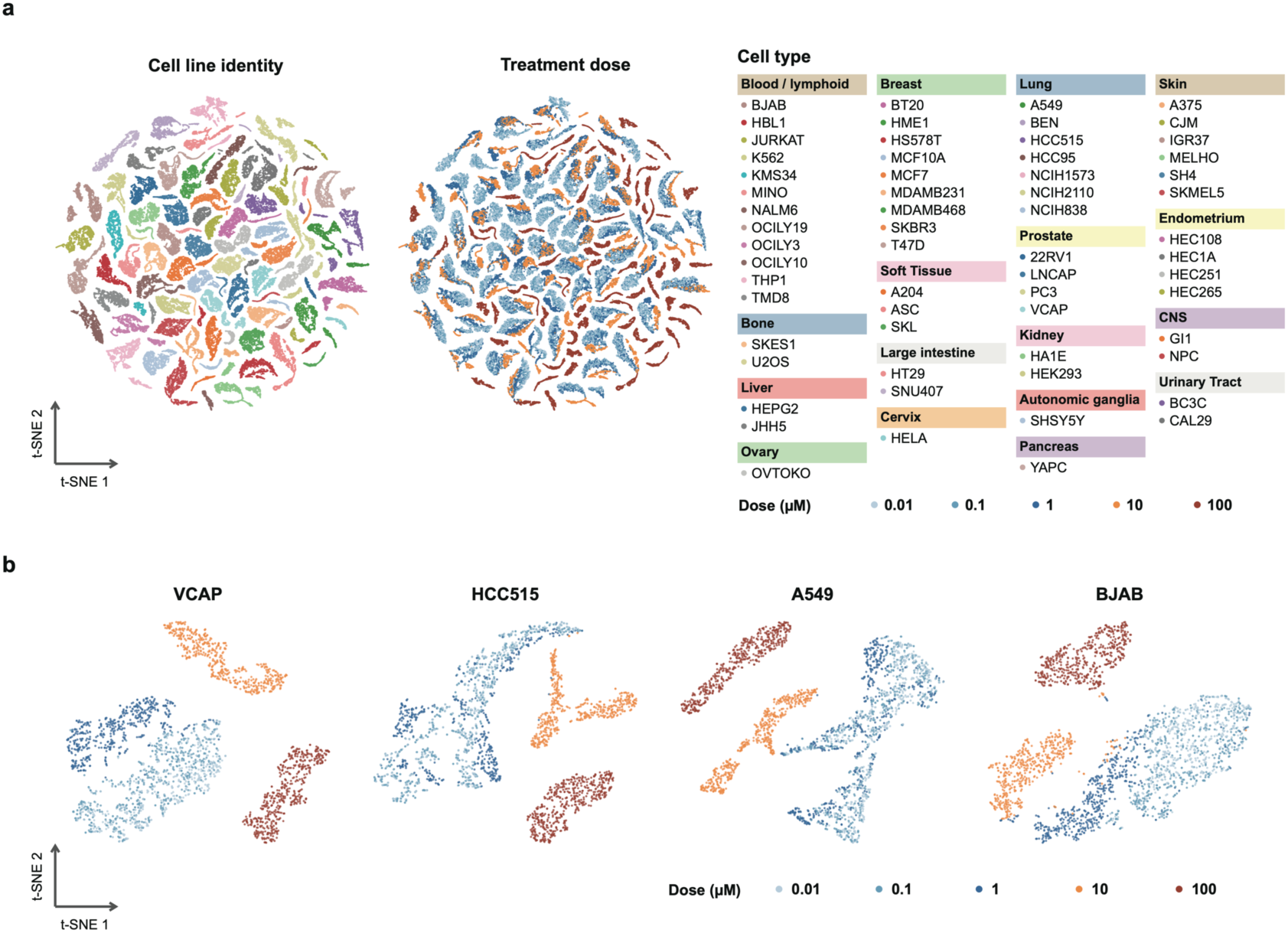
Dimensionality reduction of compound induced transcriptional profiles across cancer cell lines and doses. (a) t-SNE visualization of transcriptional profiles from all tested cell lines. Points are colored by cell line identity in the left panel and by compound dose in the right panel. The distribution indicates that transcriptional profiles cluster according to cell identity, while dose-dependent effects are also observed across the global embedding. Cell lines are grouped and annotated by tissue or cancer type. (b) Independent t-SNE visualizations for representative individual cell lines. Each panel shows transcriptional profiles generated from 516 compounds tested at five doses, with points colored by dose. Dose-dependent separation is observed within individual cell lines.

**Figure S2.**
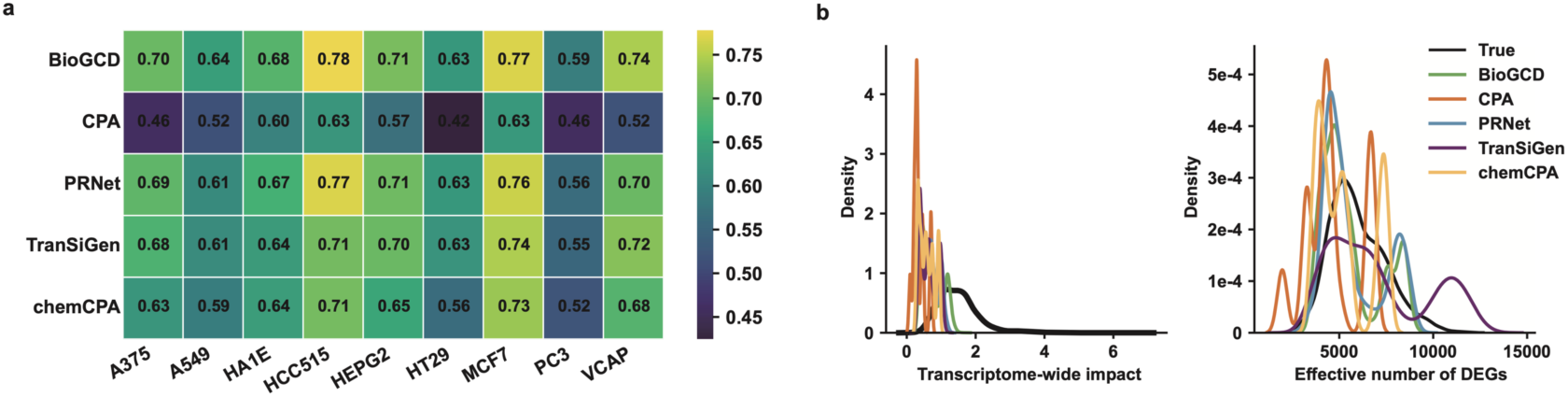
Model comparison for recovering transcriptome-wide perturbation effects. (a) Heatmap of model performance across cell lines. Higher values indicate better agreement with true perturbation responses. BioGCD shows the most consistent performance across cell lines, followed by PRNet, whereas CPA performs relatively weaker. (b) Density plots of true and predicted perturbation metrics. The left panel shows transcriptome-wide impact, and the right panel shows the effective number of DEGs. Compared with the true data, most models compress the dynamic range of perturbation effects, especially for transcriptome-wide impact. BioGCD and PRNet better approximate the true distributions.

**Figure S3.**
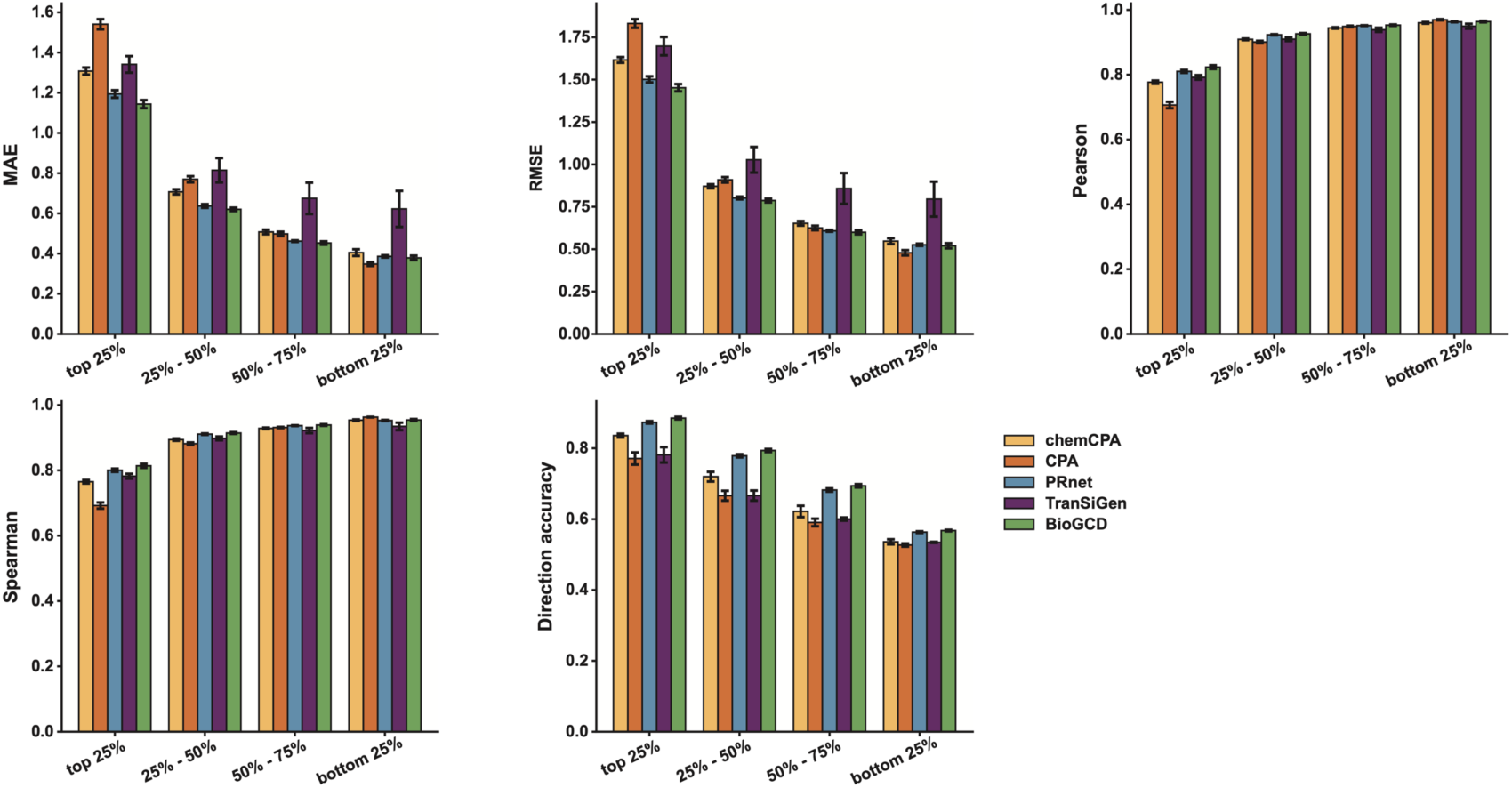
Model performance across gene quantiles. Genes were grouped into quartiles by the magnitude of observed expression changes relative to controls. Strongly perturbed genes showed higher prediction errors but greater directional accuracy, while correlations remained high across quartiles. BioGCD showed stable and competitive performance, followed by PRnet. The other models were more variable.

**Figure S4.**
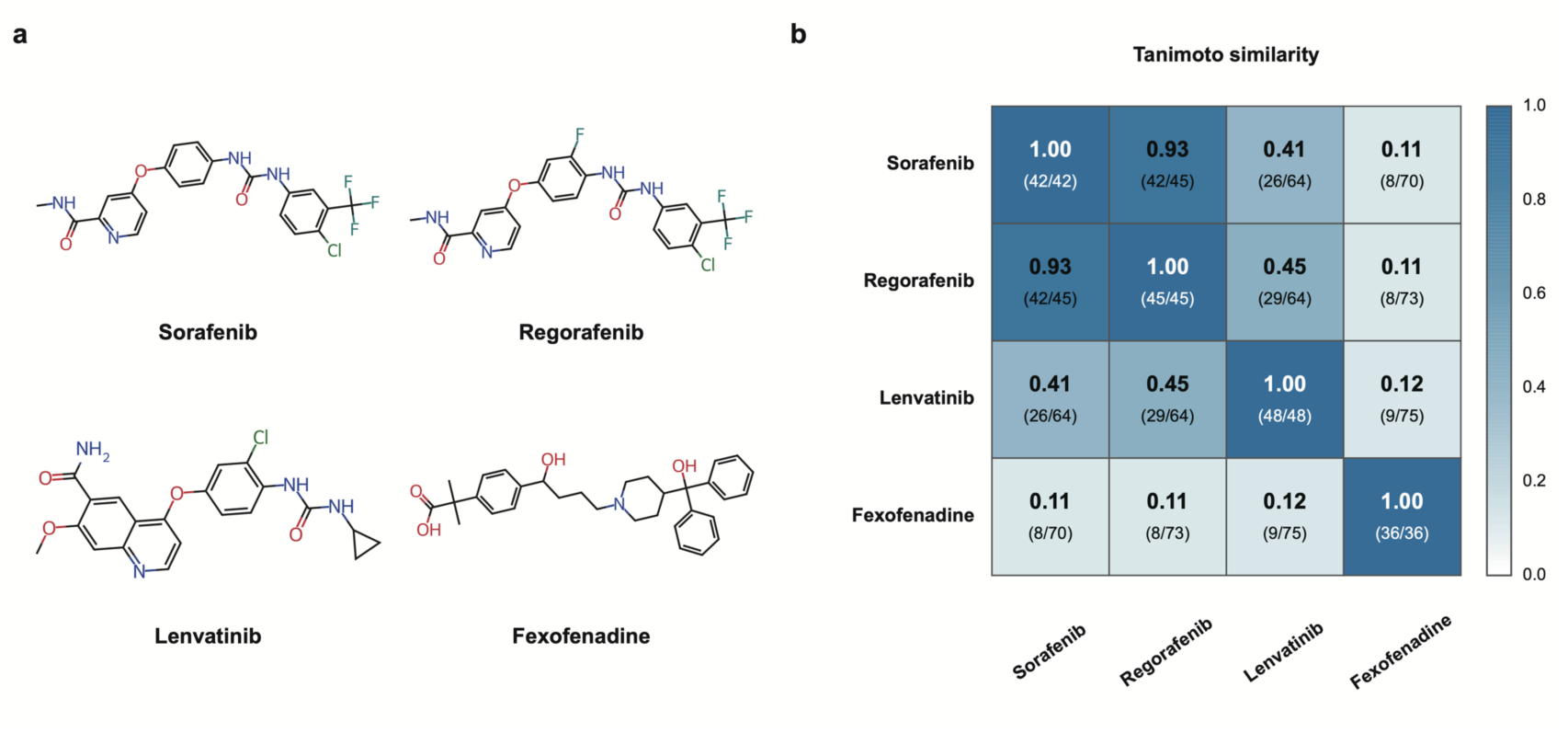
Molecular similarity. (a) Chemical structures of the selected effective drugs. Drugs with similar active fingerprint patterns, such as Sorafenib and Regorafenib, also show similar molecular structures. (b) Heatmap of Tanimoto similarity based on active molecular fingerprint bins. Sorafenib and Regorafenib show the highest similarity, while Fexofenadine is clearly distinct from the other drugs.

**Figure S5.**
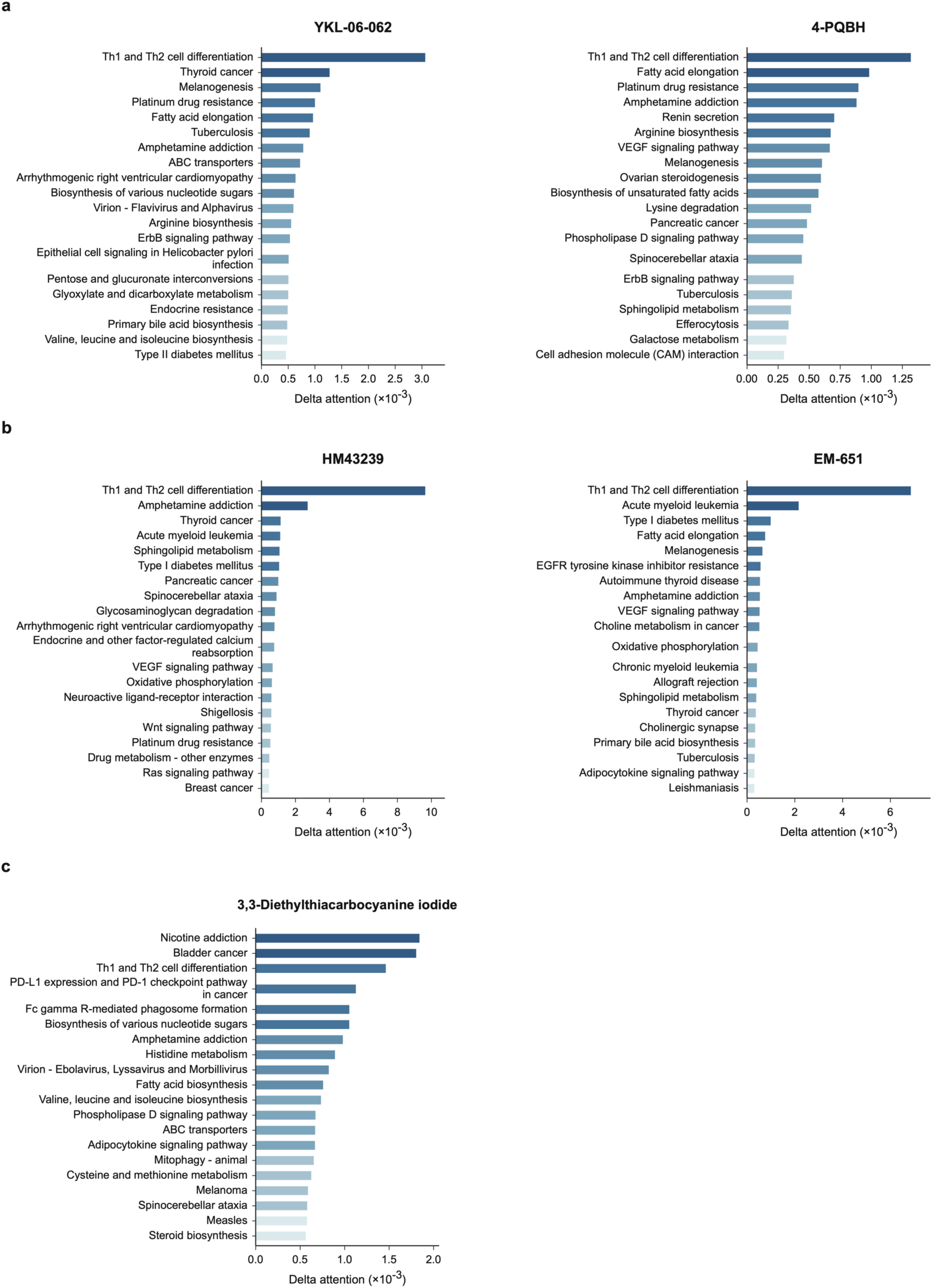
Model inferred pathway-level attention changes across compound treated cancer cell lines. Top 20 pathways ranked by increased delta attention scores for (a) YKL-06-062 and 4-PQBH in melanoma cells, (b) HM43239 and EM-651 in liver cancer cells, and (c) 3,3-diethylthiacarbocyanine iodide in breast cancer cells. Higher values indicate greater model assigned relevance under compound treatment.

**Figure S6.**
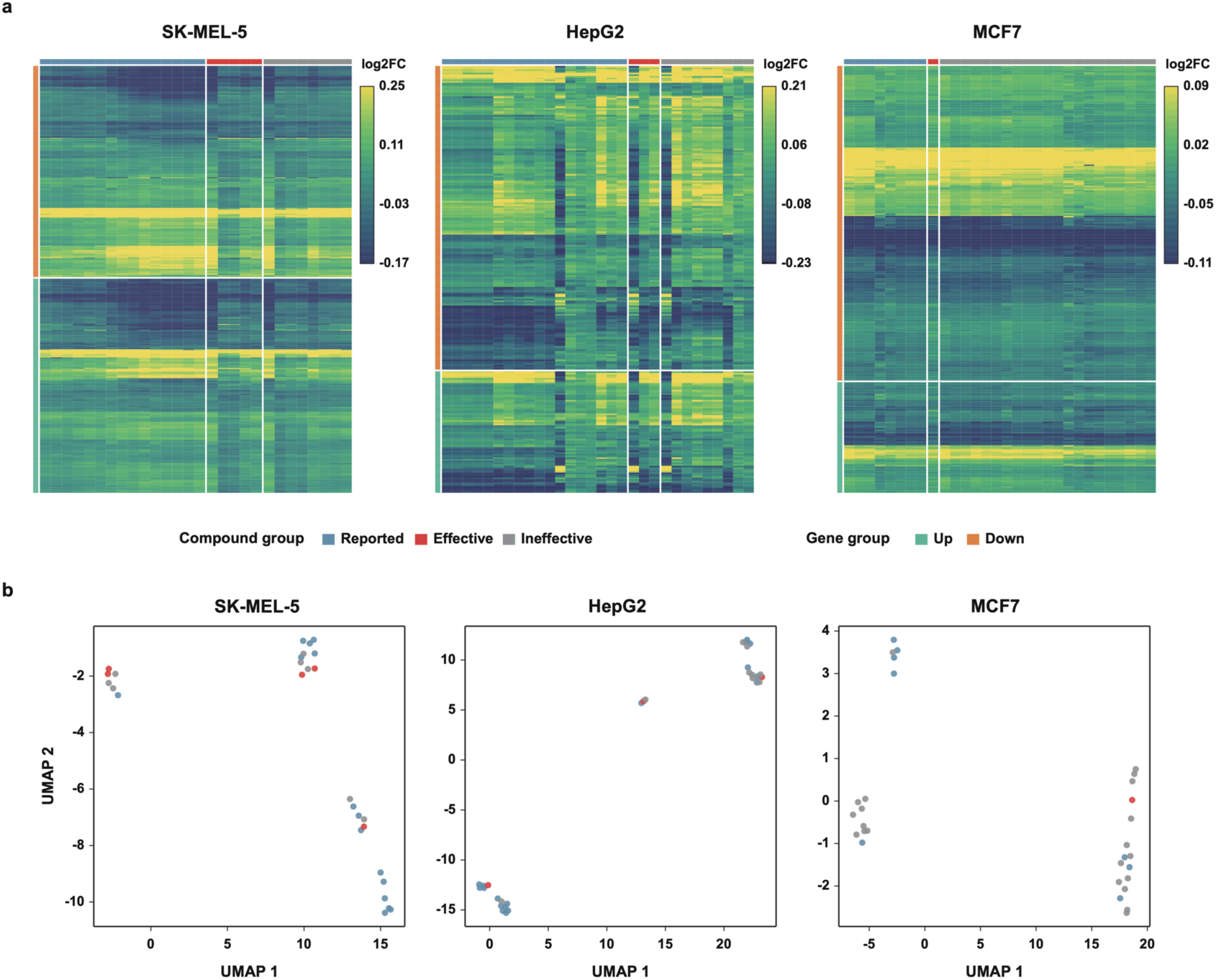
Model derived transcriptomic response patterns across compound-treated cancer cell lines. (a) Heatmaps showing log2 fold-change profiles of differentially regulated genes across predicted compound-treated SK-MEL-5, HepG2, and MCF7 cells. Rows represent genes grouped by up- or down-regulation, and columns represent compounds annotated as reported, effective, or ineffective. (b) UMAP visualization of compound-induced transcriptomic profiles in SK-MEL-5, HepG2, and MCF7 cells. Points are colored by compound group, including reported, effective, and ineffective compounds

**Figure S7.**
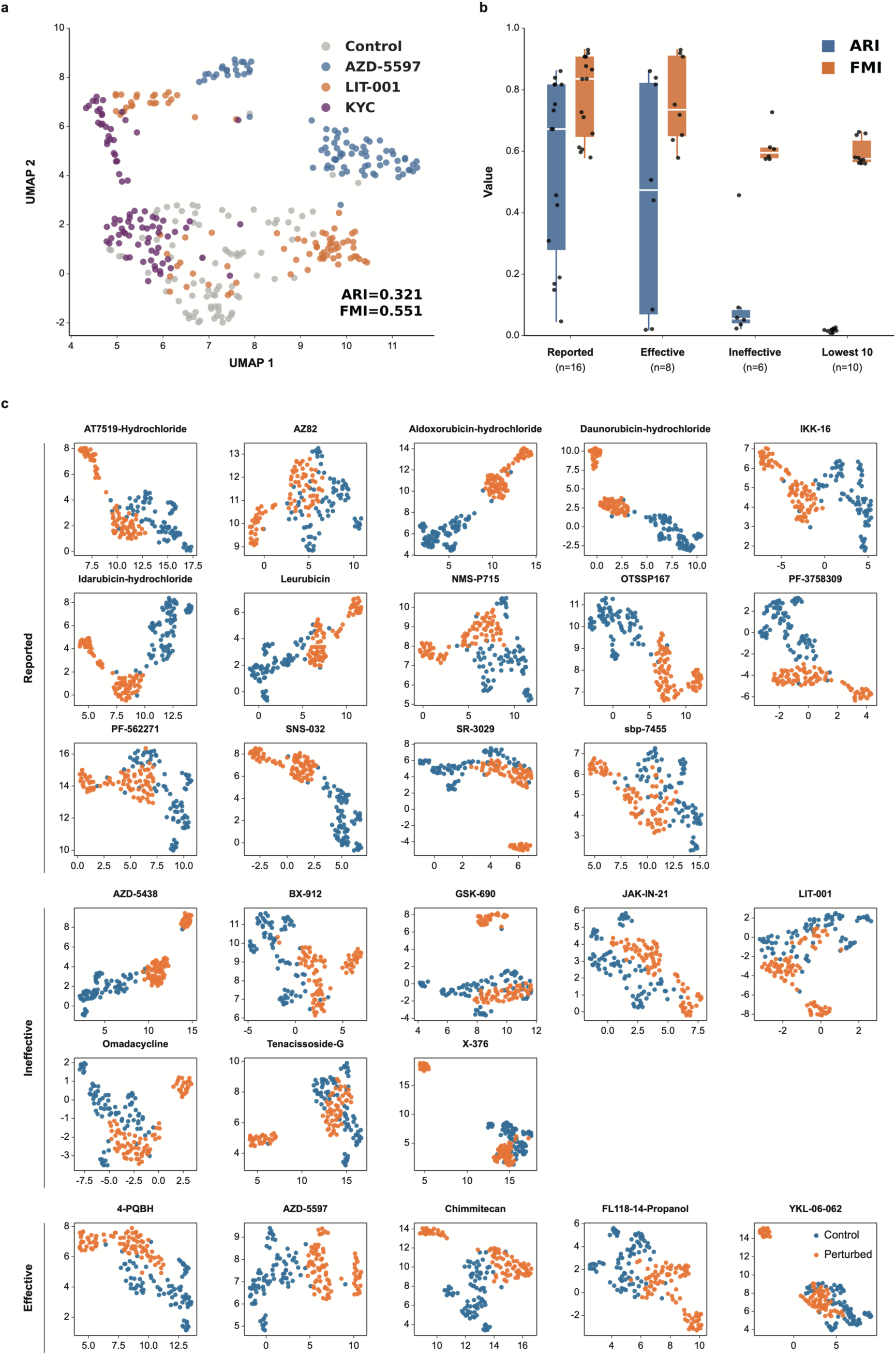
UMAP visualization of dimension-reduced profiles across compound groups. (a) Representative UMAP of control profiles and AZD-5597, LIT-001 and KYC perturbations. (b) ARI and FMI distributions across the indicated compound groups. (c) UMAP projections of dimension-reduced cellular profiles for compounds categorized as Reported, Ineffective, or Effective. Each panel shows one compound, and points are colored by sample condition, with blue indicating controls and orange indicating compound perturbed samples.

**Figure S8.**
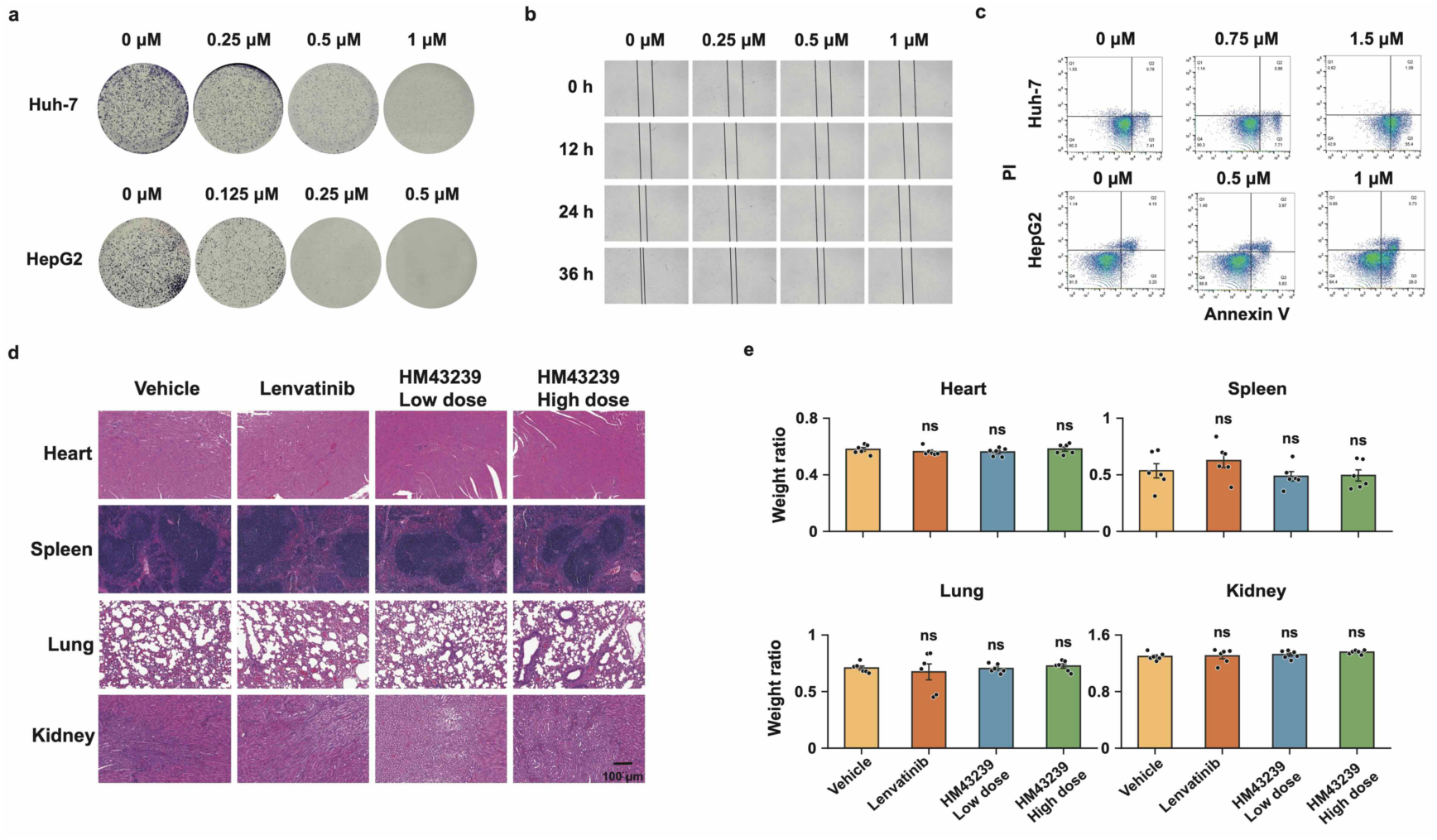
HM43239 activity in hepatocellular carcinoma cells and assessment of organ toxicity in mice. (a) Colony formation images of Huh-7 and HepG2 cells treated with HM43239 at the indicated concentrations. (b) Wound healing images of Huh-7 cells treated with HM43239 at the indicated concentrations and time points. (c) Annexin V/propidium iodide flow cytometry plots of Huh-7 and HepG2 cells treated with HM43239. (d) H&E-stained images of the heart, spleen, lung, and kidney from mice treated with vehicle, lenvatinib, or HM43239 at low or high doses. Scale bar, 100 μm. (e) Organ weight ratios for the heart, spleen, lung, and kidney across treatment groups. Data comparisons were performed using one-way ANOVA followed by Dunnett’s test against the vehicle group. ns, not significant (p ≥ 0.05).

**Figure S9.**
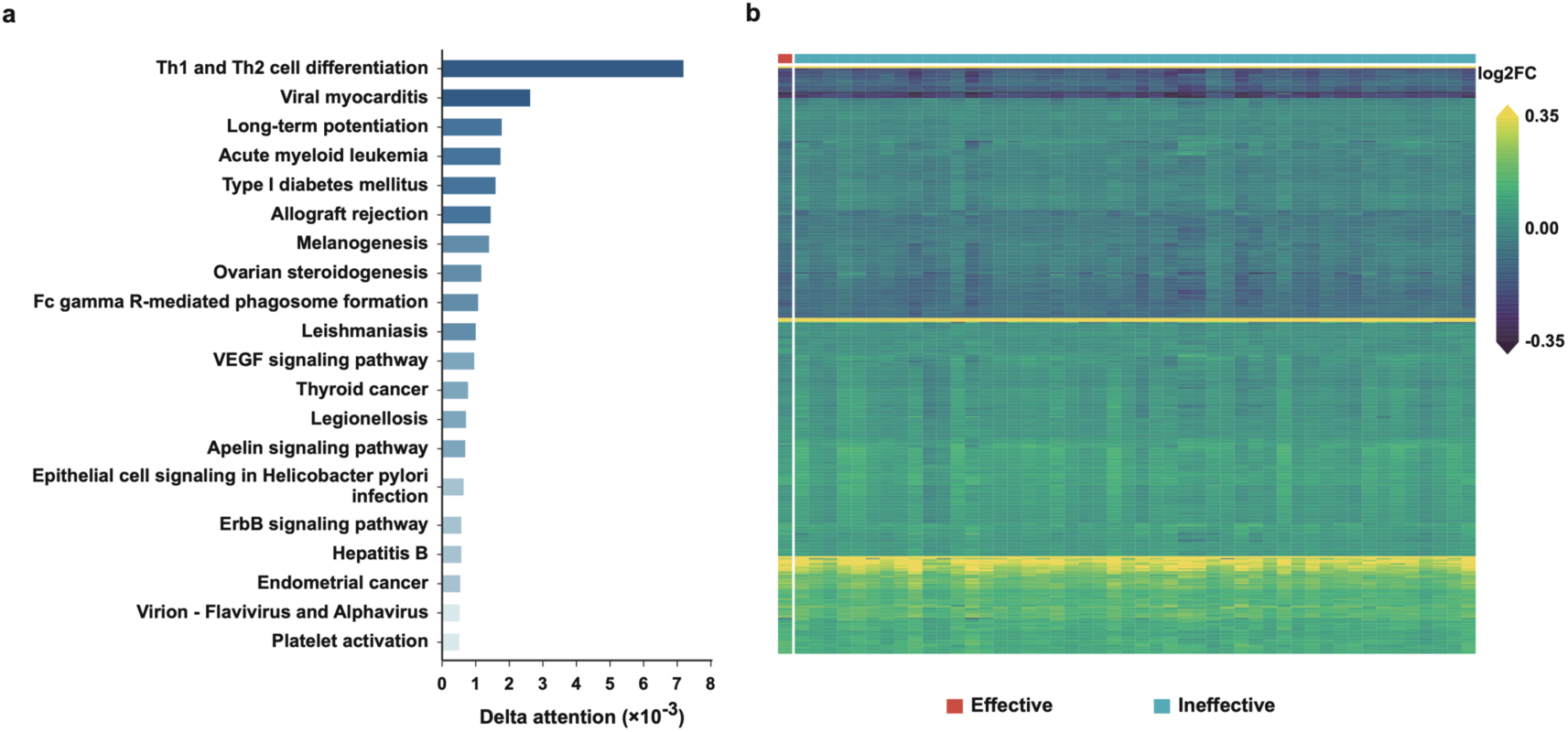
Pathway and gene expression interpretation of novel candidate compound. (A) Top KEGG pathways associated with compound Z6412951658, ranked by delta attention score. (B) Heatmap of predicted log2 fold-change values for melanoma disease-associated genes across candidate compounds. Candidate compounds are annotated as effective or ineffective, and colors indicate predicted expression changes.

**Figure S10.**
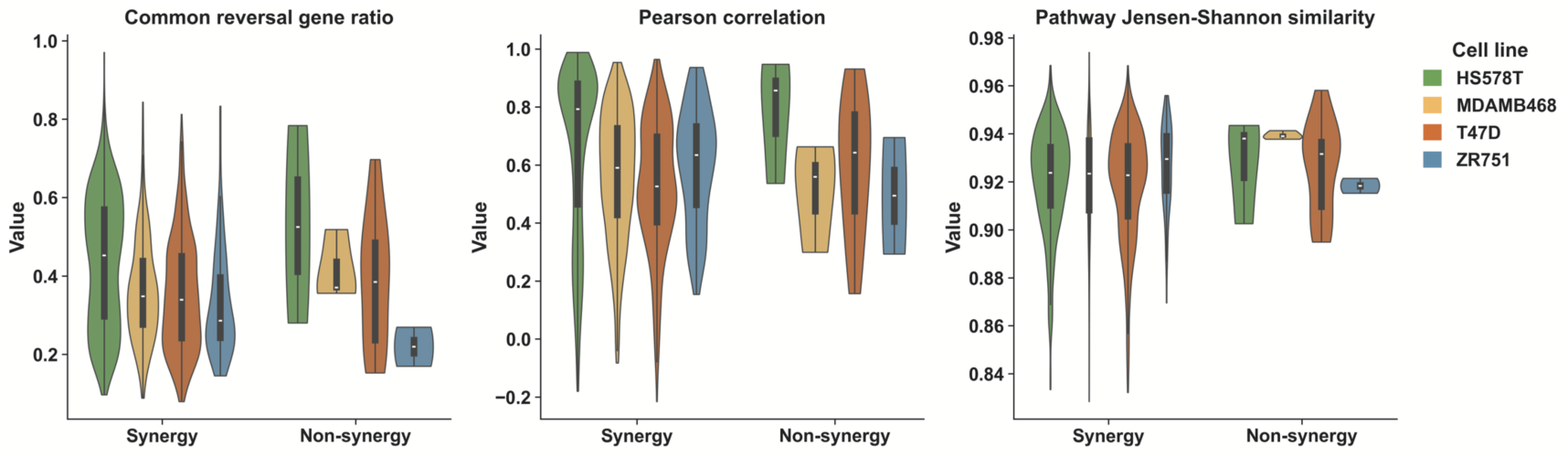
Comparison of gene-level and pathway-level features between synergistic and non-synergistic compound pairs. Violin plots showing the distributions of the common reversal gene ratio, Pearson correlation, and pathway Jensen-Shannon similarity for synergistic and non-synergistic compound pairs across four breast cancer cell lines.

**Figure S11.**
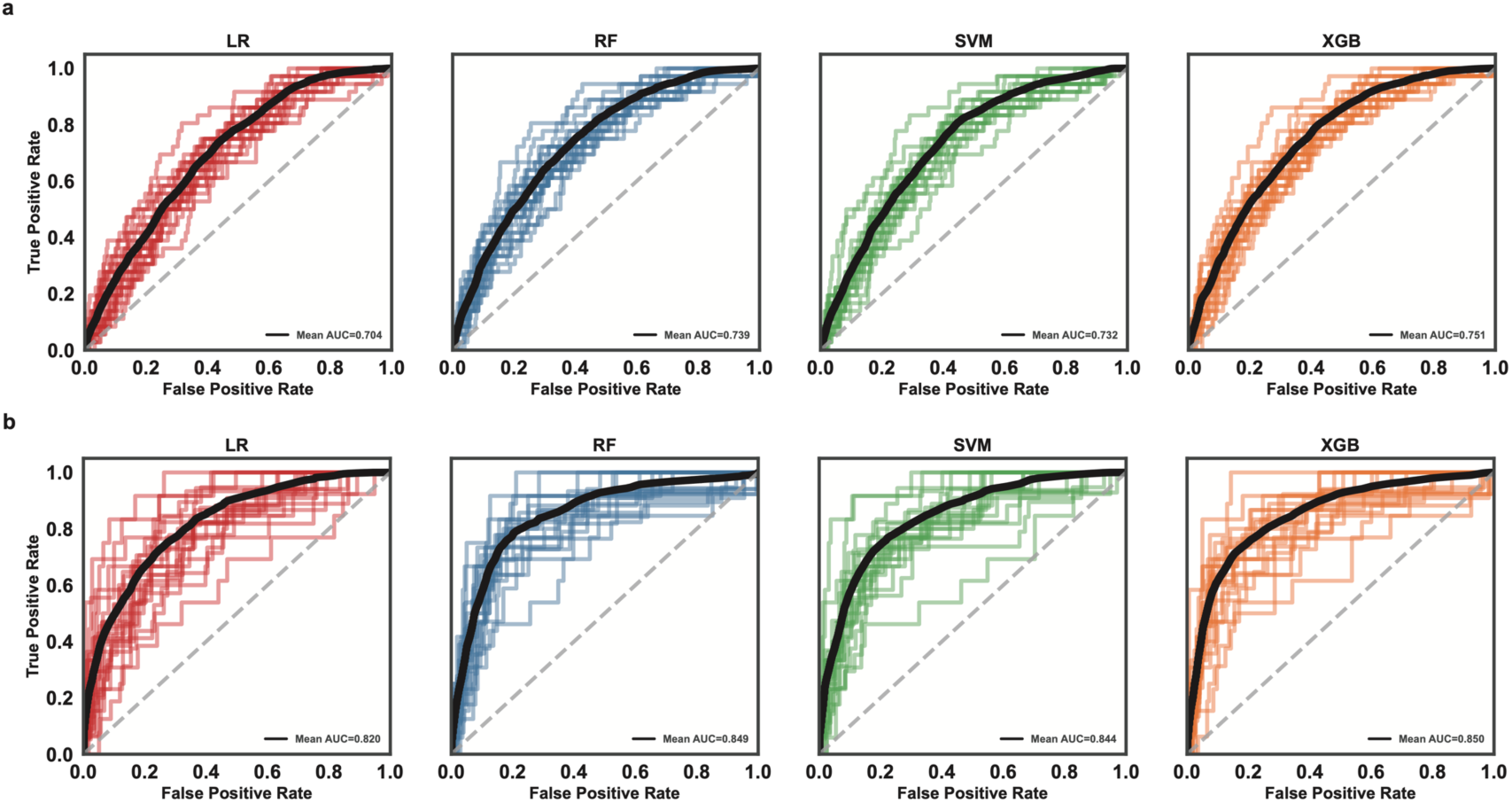
ROC-based performance comparison of machine learning models for compound synergy prediction. (a) Model performance based on three melanoma cell lines. (b) Model performance based on the SK-MEL-5 cell line. ROC curves are shown for logistic regression (LR), random forest (RF), support vector machine (SVM), and XGBoost (XGB) models across repeated cross-validation runs. Thin colored lines indicate individual runs, and black lines indicate mean ROC curves.

